# Synthesis of USC-093 and comparison with its promoiety enantiomer USC-093D against adenovirus in vitro and in a Syrian hamster model

**DOI:** 10.1101/2024.11.01.621456

**Authors:** Jiayi Yang, Samantha B. Riemann, Jinglei Lyu, Sudi Feng, Yang Bi, Nicholas A. Lentini, Inah Kang, Boris A. Kashemirov, Caroll B. Hartline, Scott H. James, Ann E. Tollefson, Anna Cline-Smith, Karoly Toth, Charles E. McKenna

## Abstract

Adenovirus infections of immunocompromised humans are a significant source of morbidity and mortality. At present, no drug has been approved by FDA for the treatment of adenovirus infections. A current treatment of such infections is off-label use of an antiviral acyclic nucleotide phosphonate, cidofovir (CDV, (*S*)-HPMPC), which requires i.v. administration and has dose-limiting kidney toxicity. We recently reported that USC-093, a homoserinamide analogue of the tyrosinamide (*S*)- HPMPA prodrug USC-087, was orally effective at a 10 mg/kg against disseminated human adenovirus infection (HAdV-C6) in a Syrian hamster model, although their efficacy was marginal after respiratory infection. Neither prodrug manifested GI toxicity. Unlike USC-087, USC-093 showed no significant nephrotoxicity at the effective dose. Here, we describe in detail the synthesis of USC-093 and also its D-homoserinamide analogue, USC-093D, in four steps (20-40% overall yield) starting from Boc-protected L-homoserine or D-homoserine lactone, respectively. The two stereoisomeric prodrugs had EC_50_ 30-70 nM vs. Ad5 or 1-6 nM vs. Ad6 in HFF cells, with USC-093D giving the lower values. The prodrugs were 30-59x more potent vs. Ad5 and 82-332x more potent than Ad6 relative to the positive control, CDV. To ascertain whether D-chirality in the promoiety could enhance the performance of the prodrug in vivo, USC-093D and USC-093 were compared in the Syrian hamster model (treated from day 1 q.d at an experimentally determined maximum tolerated oral dose of 20 mg/kg)). In this study, the hamsters were instilled i.n. with vehicle or 4X10^10^ PFU/kg of HAdV-C6 to promote lung infection. Oral valganciclovir (VGCV) at 200 mg/kg b.i.d. was used as the positive control. The body weights were recorded daily, and at 3 days post challenge, gross pathological observation was performed. Lung samples were collected, and the virus burden was determined by TCID_50_ assay. The results show that altering homoserine stereochemistry did not markedly improve the efficacy of the orally administered prodrug, consistent with the premise that its mechanism of transport is likely not dependent on stereoselective pathways, such as hPEPT1-mediated uptake.

## 1. INTRODUCTION

Human adenoviruses (HAdV) are classified into seven species and over 100 types. They are trophic to the epithelium; types belonging to species B, C, and E cause respiratory illnesses, those in species D cause keratoconjunctivitis, while the two types of species F cause gastrointestinal infections (reviewed in [1]). Most HAdV infections occur in children who lack immunity to the virus; HAdV was identified as the causative agent in 20% of infantile pneumonia cases resulting in hospitalization [2]. In immunocompetent adults, HAdV infections generally result in mild infections that resolve without serious sequelae and generate life-long immunity to the infecting type [3]. However, there are several exceptions to this. Epidemic keratoconjunctivitis (caused by species D HAdV) can result in permanent eye damage [4], and HAdVs (mostly types HAdV-E4 and HAdV-B7) also account for the majority of acute respiratory cases among military recruits [3]. Lately, emerging HAdV types have caused sporadic infections and local outbreaks resulting in fatalities [3, 5]. HAdV may also be implicated in an upsurge of pediatric hepatitis cases with unknown etiology [6]. Still, the most severely affected patient population is that of the immunocompromised. As more and more people receive life-saving transplantation treatment, there is a growing population of severely immunocompromised patients who are very susceptible to opportunistic infections. Especially vulnerable are pediatric allogeneic hematopoietic stem cell recipients, for whom lymphocytes are depleted during conditioning and during the engraftment period [7]. This protracted, severe immunosuppression regimen makes these patients susceptible to dsDNA virus infections that would be otherwise eliminated by a healthy immune system [8]. HAdV are important pathogens in this population, with up to 40% incidence rate and a 10% case fatality rate. The fatality rate may reach 80% in the case of disseminated disease [9, 10]. The source of HAdV infection can be the donor tissue or it can be community acquired, as well persistent infection of the recipient may be reactivated during immunosuppression [11]. Transplant patients are routinely monitored for viral infections; detecting HAdV DNA in the blood and/or stool is an important biomarker that precedes disease symptoms and allows medical intervention [11]. Based on the epidemiology of HAdV infections, there is a clear need for effective countermeasures to HAdV infection.

To date, there is no drug specifically approved by the FDA to treat HAdV infections. While there are ongoing efforts to reconstitute the lymphocyte compartment to provide immune protection against the infection (reviewed in [12]), the main thrust of research is to develop a small-molecule antiviral agent. Presently, most clinicians resort to the off-label use of cidofovir (CDV), a drug approved to treat CMV retinitis, to combat adenovirus infections [13]. CDV is an acyclic nucleotide phosphonate that is phosphorylated to the active diphosphate form by cellular kinases and acts as a chain terminator and competitive inhibitor of the HAdV polymerase during viral DNA replication [14]. However, CDV is a substrate for the anionic transporter, and this, combined with the long plasma half-life of the molecule, causes severe nephrotoxicity [15]. Brincidofovir (BCV), a lipid-linked derivative of CDV, demonstrated excellent anti-adenoviral activity in vitro and in vivo, and showed improved oral bioavailability and reduced kidney toxicity [16-18]. BCV reached Phase III clinical trials for treatment of HAdV infections, but the oral administration had to be abandoned because of enterotoxicity that could not be distinguished from graft-versus-host disease [19-21]. Presently, BCV is being developed as an intravenously (i.v.) administered antiviral drug [22]. While this approach is valid for hospitalized patients, oral therapy still may be necessary for home care.

We recently reported that USC-093 [23], a homoserinamide analogue of the tyrosinamide (*S*)-HPMPA prodrug USC-087 (**Figure 1**), was orally effective at a 10 mg/kg against disseminated human adenovirus infection (HAdV-C6) in a Syrian hamster model, although their efficacy was marginal after respiratory infection. Neither prodrug manifested GI toxicity. Unlike USC-087, USC-093 showed no significant nephrotoxicity at the effective dose. Here, we describe in detail the synthesis of USC-093 and also its novel D-homoserinamide analogue USC-093D in four steps (20-40% overall yield) starting from Boc-protected L-homoserine or D-homoserine lactone, respectively. To ascertain whether D-chirality in the promoiety could enhance the performance of the prodrug in vivo, USC-093D and USC-093 were compared in the Syrian hamster model using i.n. instillation of HAdV-C6 to promote lung infection. The results are used to assess the possibility that these prodrugs utilize a mechanism of transport dependent on stereoselective pathways, such as hPEPT1-mediated uptake.

**Figure 1.**
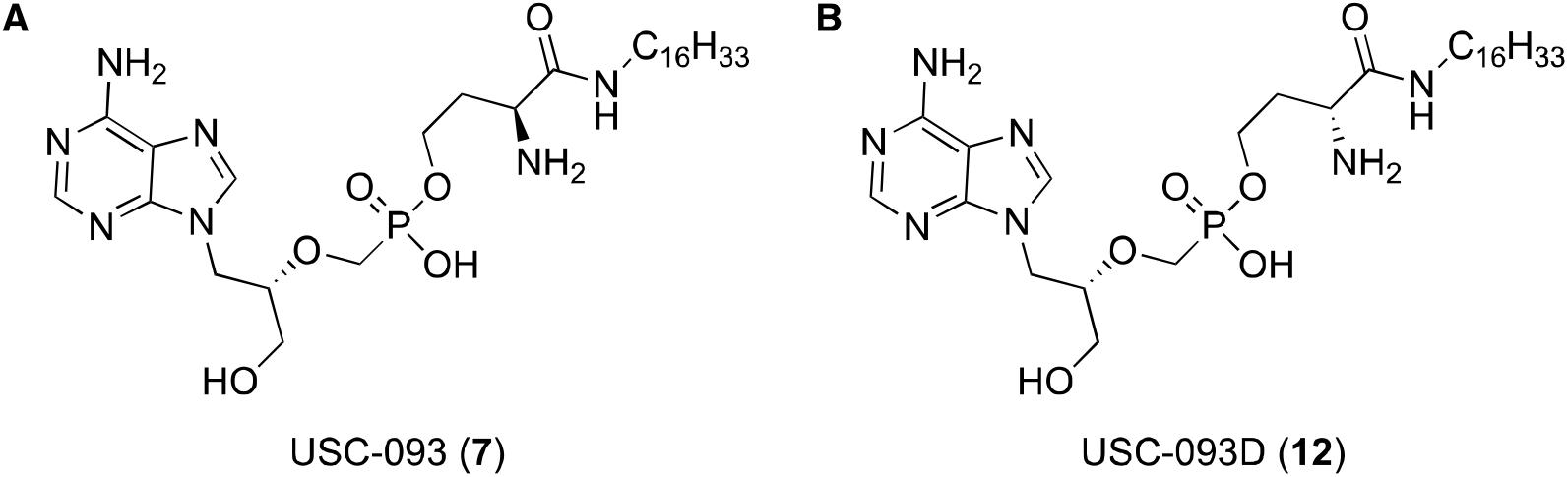
USC-093 (7) and USC-093D (12); these HPMPA prodrugs differ only in the chirality of the α-carbon of the homoserine moiety.

**Figure 2.**
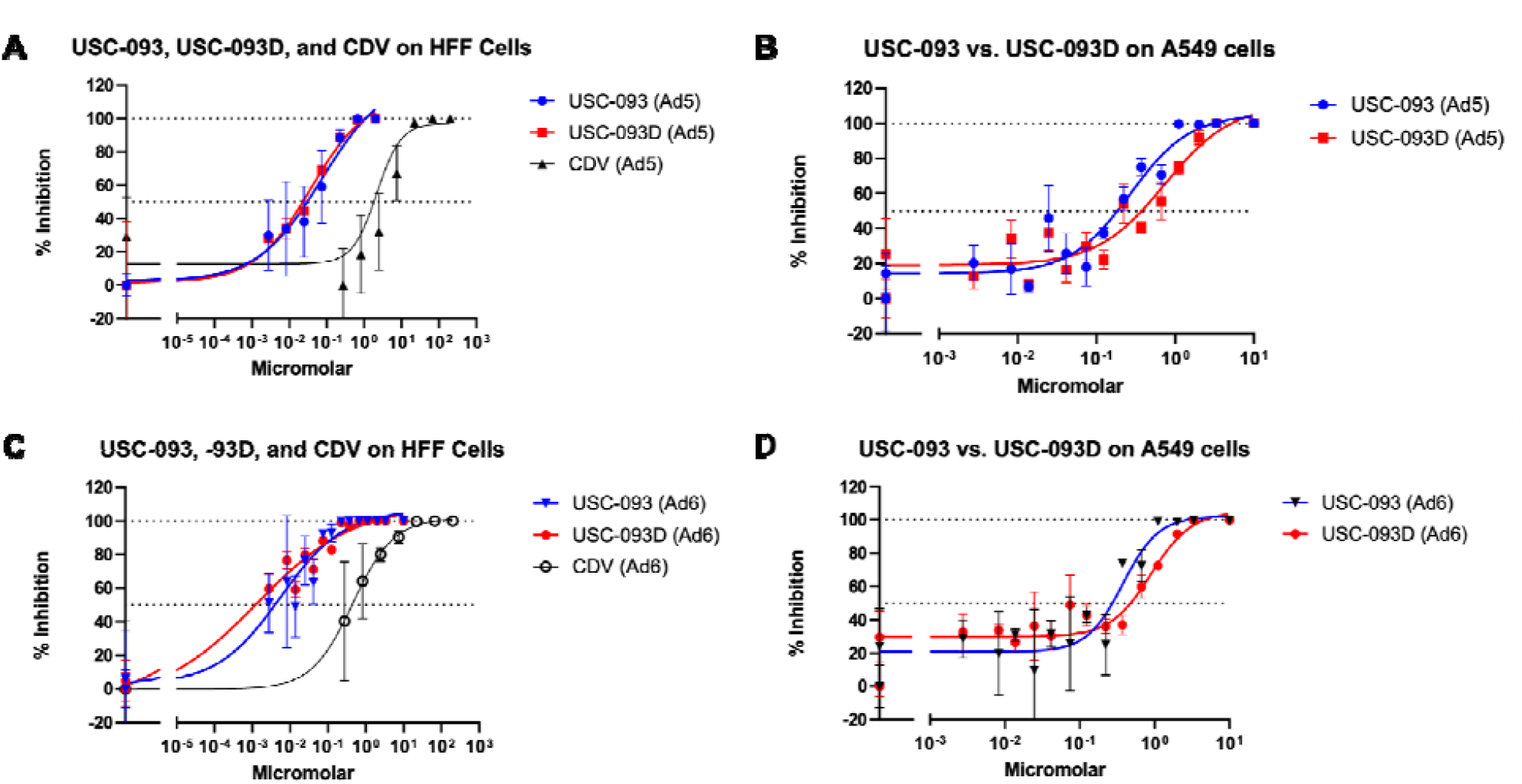
Inhibition by USC-093, USC-093D of Ad5 and Ad6 in cell culture. A) Evaluation of USC-093, USC-093D, and CDV (positive control) against Ad5 in HFF cells. B) Evaluation of USC-093 and USC-093D against Ad5 in A549 cells. C) Evaluation of USC-093, USC-093D, and CDV against Ad6 in HFF cells. D) Evaluation of USC-093 and USC-093D against Ad6 in A549 cells.

## 2. MATERIALS AND METHODS

### 2.1 Chemistry

If not stated otherwise, all reagents and solvents were purchased from Sapala Organics, VWR, Sigma Aldrich, Fisher, TCI, or Oakwood and used without further purification. Products were purified by flash column chromatography on Silica RediSep Rf prepacked columns of appropriate size using a Teledyne ISCO CombiFlash Rf+ Lumen system equipped with UV-Vis and evaporate light scattering detectors. NMR spectra were obtained on either Varian VNMRS-500 or Varian Mercury 400 2-channel spectrometers with CD_2_HOD as internal standard for ^1^H and 85% H_3_PO_4_ as external standard for ^31^P, all ^31^P NMR were proton-decoupled, and all NMR samples used CD_3_OD as solvent unless specified. MS was performed using the Thermo-Finnigan LCQ Deca XP Max system or Advion Interchim Scientific expression compact mass spectrometer (CMS) with ESI detection. Prodrugs were further characterized by UV/Vis spectroscopy (Beckman Coulter DU 800 spectrophotometer) combustion elemental analysis (Galbraith Laboratories) and by mass spectrometry (MS).

#### 2.1.1 Synthesis of prodrug 7 (USC-093)

##### *tert*-Butyl (*S)-(1-(hexad*ecylamino)-4-hydroxy-1-oxobutan-2-yl)carbamate, 3

*tert* -Butyl (*S*)-homoserine lactone **1** (1.5 g, 1 eq, 7.45 mmol) and hexadecane-1-amine **2** (2.2 g, 1.2 eq, 8.95 mmol) were dissolved in 25 mL of THF. The mixture was heated at 60 °C, stirred for 2 d, and volatiles were evaporated under reduced pressure. The residue was then subjected to flash chromatography on a silica column (Teledyne ISCO 40 g Flash Column) eluted with a gradient of hexanes and EtOAc from 0 – 100% to yield the promoiety 3 (2.87 g, 87%) at 50% EtOAc (**Figure S1**). ^1^H NMR (400 MHz, CDCl_3_) δ 6.02 – 5.88 (m, 1H), 5.56 – 5.44 (m, 1H), 4.32 – 4.22 (s, 1H), 3.76 – 3.67 (m, 2H), 3.26 (q, *J* = 6.7 Hz, 2H), 2.01 – 1.93 (m, 1H), 1.81 – 1.70 (m, 1H), 1.45 (d, *J* = 0.8 Hz, 9H), 1.34 – 1.21 (m, 26H), 0.88 (t, *J* = 6.7 Hz, 3H) (**Figure S2**).

##### *tert*-Butyl ((2S)-(4-(((5R)-5-(6-amino-9H-purin-9-yl)-2-oxido-1,4,2-dioxaphosphinan-2-yl)oxy)-1-(hexadecylamino)-1-oxobutan-2-yl)carbamate, 5

(*S*)-HPMPA **4** (791 mg, 1 eq, 2.61 mmol), 1.5 g of promoiety **3** (1.3 eq, 3.39 mmol) and 3.8 g of PyBOP (2.80 eq, 7.30 mmol) were mixed with 25 mL of freshly distilled DMF and 8.6 mL of freshly distilled DIEA (19 eq, 49.5 mmol). The mixture was stirred at 40°C for 2 h while the reaction was monitored by ^31^P NMR to observe completion. After removal of volatiles under reduced pressure, the residue was semi-purified via automated flash chromatography on a silica column (Teledyne ISCO 24 g flash column) eluted with a gradient of DCM and MeOH from 0 – 30% to provide the oxaphosphinane diastereomers **5** (1.83 g, 99%) (Figure S3). ^1^H NMR (400 MHz) δ 8.26 (s, 1H), 8.14 (d, *J* = 7.7 Hz, 1H), 4.47 – 4.39 (m, 2H), 4.38 – 4.15 (m, 6H), 3.26 – 3.15 (m, 2H), 2.28 – 2.10 (m, 1H), 2.01 – 1.86 (m, 1H), 1.49 (d, *J* = 1.4 Hz, 6H), 1.45 (s, 3H), 1.41 – 1.25 (m, 30H), 0.93 (t, *J* = 6.5 Hz, 3H) (**Figure S4**). ^31^P NMR (162 MHz, CDCl_3_) δ 12.87, 11.34 (**Figure S5**).

##### *tert*-Butyl (2S)-(4-(((((*R)-1-(6-amin*o-9*H*-purin-9-yl)-2-hydroxyethoxy)methyl)(hydroxy)phosphoryl)-oxy)-1-(hexadecylamino)-1-oxobutan-2-yl)carbamate, 6

Oxaphosphinanes **5** (1.4 g, 1 eq, 1.99 mmol) were combined with a mixture of 53 mL of saturated aq. NH_4_OH (14.8 M, 394 eq, 784 mmol) and 143 mL of MeCN. The resultant mixture was heated at 45 °C and stirred for 6 d while the reaction was monitored by ^31^P NMR. Volatiles were then evaporated under reduced pressure and the residue was subjected to automated flash chromatography using a silica column eluted with a gradient of DCM and MeOH from 0 – 100% to isolate phosphonate **6** (890 mg, 62%) at 30% MeOH (**Figure S6**). H NMR (500 MHz) δ 8.29 (s, 1H), 8.23 (s, 1H), 4.48 (dd, *J* = 14.5, 3.9 Hz, 1H), 4.39 (dd, *J* = 14.2, 6.8 Hz, 1H), 4.19 – 4.13 (m, 1H), 3.98 – 3.88 (m, 2H), 3.84 – 3.57 (m, 4H), 3.51 (dd, *J* = 12.6, 4.4 Hz, 1H), 3.25 – 3.16 (m, *J* = 6.6 Hz, 2H), 2.07 – 1.96 (m, 1H), 1.95 – 1.88 (m, 1H), 1.56 – 1.49 (m, 2H), 1.42 (s, 9H), 1.34 – 1.26 (m, 26H), 0.92 (t, *J* = 6.9 Hz, 3H) (**Figure S7**). ^31^P NMR (202 MHz): δ 16.21 (**Figure S8**).

##### (*S*)-3-amino-4-(hexadecylamino)-4-oxobutyl hydrogen (((R)-1-(6-amino-9H-purin-9-yl)-2-hydroxyethoxy)methyl)phosphonate, 7 (USC-093)

Phosphonate **6** (382 mg, 1 eq, 0.52 mmol) was dissolved in 240 mL of THF at 0 °C and the mixture was stirred. Saturated hydrochloric acid (12.1 mL, 38% by weight, 0.044 eq, 0.15 mmol) was added dropwise while stirring. The progress of the reaction was monitored by mass spectrometry for 24 hours. After 24 hours, the volatiles were evaporated under reduced pressure and the residue subjected to automated flash chromatography using a silica column eluted with a gradient of DCM and MeOH from 0 – 100%, the phosphonate **7** was eluted at 30% MeOH. The product containing fractions were combined, evaporated under reduced pressure, the material that remained was triturated with ether to afford **7** (250 mg, white powder, 76%) – USC-093 (**Figure S9**). To prepare for in vivo studies, we synthesized 1 gram total of USC-093 in four separate batches. Overall Yield 40%. ^1^H NMR (500 MHz) δ 8.26 (s, 1H), 8.23 (s, 1H), 4.51 – 4.33 (m, 2H), 4.04 – 3.88 (m, 3H), 3.82 (dq, *J* = 8.2, 4.2 Hz, 1H), 3.74 (ddd, *J* = 13.1, 7.2, 3.1 Hz, 2H), 3.64 – 3.49 (m, 2H), 3.24 (t, *J* = 7.1 Hz, 2H), 2.12 (ddt, *J* = 14.8, 7.6, 4.8 Hz, 1H), 1.97 (dddd, *J* = 14.9, 9.8, 6.2, 3.9 Hz, 1H), 1.54 (p, *J* = 7.1 Hz, 2H), 1.29 (s, 28H), 0.90 (t, *J* = 6.9 Hz, 3H) (**Figure S10**). ^31^P NMR (162 MHz): δ 17.47 (**Figure S11**). LR-MS (ESI+, *m/z*) calcd for [M+H]^−^ C_29_H_55_N_7_O_6_P^−^ : 628.39; found: 628.6 (ESI–, *m/z*) calcd for [M–H]^−^ C_29_H_53_N_7_O_6_P^−^ : 626.39; found: 626.2 (**Figure S12**). Combustion elemental analysis, %: calcd for C_29_H_54_N_7_O_6_P: C, 55.49; H, 8.67; N, 15.62 (**Figure S13**). Found: C, 55.19; H, 8.73; N, 15.08.

#### 2.1.2 Synthesis of prodrug 12 (USC-093D)

##### *tert* -Butyl (*R*)-(1-(hexadecylamino)-4-hydroxy-1-oxobutan-2-yl)carbamate, 9

Following the synthesis of **3**, tert-butyl (*R*)-homoserine lactone **8** (1.50 g), the D-homoserine analogue of **1**, was reacted with amine **2** (2.16 g) in 38 mL of THF to yield the promoiety **9** after flash chromatography as a white powder (2.80 g, 85%) (**Figure S14**). ^1^H NMR (500 MHz, CDCl_3_) δ 6.58 (t, *J* = 5.6 Hz, 1H), 5.64 (d, *J* = 7.7 Hz, 1H), 4.34 – 4.25 (m, 1H), 3.71 – 3.67 (m, 2H), 3.22 (h, *J* = 6.7 Hz, 2H), 2.03 – 1.92 (m, 1H), 1.79 – 1.68 (m, 1H), 1.43 (s, 9H), 1.32 – 1.20 (m, 26H), 0.87 (t, *J* = 6.8 Hz, 3H) (**Figure S15**).

##### *tert*-Butyl ((2R)-(4-(((5R)-5-(6-amino-9H-purin-9-yl)-2-oxido-1,4,2-dioxaphosphinan-2-yl)oxy)-1-(hexa-decylamino)-1-oxobutan-2-yl)carbamate, 10

Following the synthesis of **5**, (*S*)-HPMPA **4** (1.32 g, 1 eq, 4.34 mmol), promoiety **9** (2.50 g, 1.3 eq, 5.65 mmol) and PyBOP (9.0 g, 4 eq, 17.4 mmol) in 42 mL DMF were reacted with DIEA (14.4 mL, 19 eq, 82.5 mmol) to provide oxaphosphinane diastereomers **10** (2.38 g, 80%). The product was purified in six different batches by flash chromatography (**Figure S16**). ^1^H NMR (600 MHz, CDCl_3_) δ 8.22 (s, 1H), 7.81 (d, *J* = 9.5 Hz, 1H), 4.48 – 3.97 (m, 10H), 3.80 (d, *J* = 12.4 Hz, 1H), 3.15 (d, *J* = 35.0 Hz, 2H), 2.06 (s, 3H), 1.38 – 1.30 (m, 9H), 1.16 (q, *J* = 12.4 Hz, 26H), 0.78 (t, *J* = 9.0 Hz, 3H) (**Figure S17**). ^31^P NMR (243 MHz, CDCl_3_) δ 13.01, 10.96 (**Figure S18**).

##### *tert*-Butyl (2R)-(4-(((((*R)-1-(6-amin*o-9*H-purin-9-yl*)-2-hydroxyethoxy)methyl)(hydroxy)phosphoryl)-oxy)1-(hexadecylamino)-1-oxobutan-2-yl)carbamate, 11

Following the synthesis of **6**, oxaphosphinanes **10** (2.38 g, 1 eq, 3.34 mmol) and 51 mL of saturated aq. NH_4_OH in 657 mL of MeCN provided phosphonate **11** after flash chromatography (1.51 g, 62%) (**Figure S19**). ^1^H NMR (400 MHz) δ 8.24 (s, 1H), 8.18 (s, 1H), 4.44 (dd, *J* = 14.5, 3.8 Hz, 1H), 4.35 (dd, *J* = 14.6, 6.6 Hz, 1H), 4.12 (t, *J* = 6.3 Hz, 1H), 3.97 – 3.81 (m, 2H), 3.81 – 3.53 (m, 4H), 3.53 – 3.43 (m, 1H), 3.22 – 3.12 (m, 2H), 2.04 – 1.93 (m, 1H), 1.93 – 1.81 (m, 1H), 1.52 – 1.44 (m, 2H), 1.39 (d, *J* = 1.8 Hz, 9H), 1.28 – 1.23 (m, 26H), 0.87 (t, *J* = 6.7 Hz, 3H) (**Figure S20**). ^31^P NMR (162 MHz) δ 16.11 (**Figure S21**).

##### (*R)-3-amino-4*-(hexadecylamino)-4-oxobutyl hydrogen ((((*S)-1-(6-amin*o-9H-purin-9-yl)-3-hydroxy-propan-2-yl)oxy)methyl)phosphonate 12 (USC-093D)

Following the synthesis of **7**, phosphonate **11** (1.51 g, 1 eq, 2.07 mmol) in 147.5 mL of THF was treated with 7.4 mL of sat. HCl to provide after flash chromatography **12** (791.10 mg, white powder, 62%) – USC-093D (**Figure S22**).

Overall Yield 22%. ^1^H NMR (600 MHz) δ 8.41 (s, 1H), 8.36 (s, 1H), 4.56 (dd, *J* = 14.5, 3.7 Hz, 1H), 4.45 (dd, *J* = 14.6, 7.1 Hz, 1H), 4.03 – 3.98 (m, 3H), 3.83 (dq, *J* = 8.1, 4.3 Hz, 1H), 3.81 – 3.76 (m, 2H), 33.65 – 3.54 (m, 2H), 3.25 (t, *J* = 7.2 Hz, 2H), 2.17 – 2.10 (m, 1H), 2.05 – 1.96 (m, 1H), 1.56 (p, *J* = 7.3 Hz, 2H), 1.41 – 1.23 (m, 26H), 0.92 (t, *J* = 7.1 Hz, 3H) (**Figure S23**). ^31^P NMR (243 MHz) δ 17.05 (**Figure S24**). ^13^C NMR (151 MHz) δ 169.98, 152.49, 150.48, 146.33, 146.28, 119.48, 81.75 (d, *J* = 11.7 Hz), 66.39 (d, *J* = 159.5 Hz), 61.79 (d, *J* = 5.8 Hz), 52.26, 45.82, 40.73, 34.01 (d, *J* = 4.5 Hz), 33.06, 30.77, 30.74, 30.73, 30.69, 30.46, 30.42, 30.33, 28.02, 23.72, 14.42 (**Figure S25**). MS (ESI–, *m/z*) calcd for [M–H]^−^ C_29_H_53_N_7_O_6_P^−^: 626.39; found: 626.2 (**Figure S26**). 93% active content by UV analysis (260 nm, HPMPA ref.).

### 2.2 Cells and viruses

Cell lines used at Saint Louis University (SLU) were A549 cells (CCL-185; ATTC) and HFF cells (gift from T. Hermiston). HAdV-C5 (plaque-purified from VR-5, Adenoid 75; ATCC) and HAdV-C6 (VR-6, Tonsil 99; ATCC) were obtained from ATCC and cultured, purified, and titered as described in [24]. HAdV-C6 was used as a challenge virus because this type replicates better than HAdV-C5 in Syrian hamsters, and the pathology induced by HAdV-C6 is more dependent on virus replication than it is for HAdV-C5 [25, 26].

For in vitro assays at University of Alabama at Birmingham (UAB), human foreskin fibroblasts (HFFs) for in vitro cytopathic effect (CPE) assays were prepared as previously described, with human foreskins obtained from the tissue procurement facility under UAB institutional review board approval [27].

### 2.3 CDV control

The CDV control for the in vitro studies was kindly provided by Gilead Sciences, Foster City, CA. CDV for in vivo studies was obtained from the National Institute of Allergy and Infectious Diseases (NIAID).

### 2.4 Antiviral assays

In vitro CPE reduction assays were conducted in triplicate independent studies in HFF cells, with test drug concentrations ranging from 0.003 to 1 µM (control drug, CDV, range 0.048-150 µM), as previously described [28, 29]. Briefly, monolayers of cells (5,000/well) were seeded in 384-well plates. Compound dilutions were prepared in duplicate wells in a series of 5-fold dilutions directly in the plates. Cell monolayers were infected with a multiplicity of infection (MOI) of approximately 0.005 PFU/cell. Cytopathology was determined by the addition of CellTiter-Glo reagent (Promega, Madison, WI) according to the manufacturer’s suggested protocol. Concentrations of test compounds sufficient to reduce CPE by 50% (EC_50_) were interpolated from the experimental data, while cytotoxicity was concurrently determined as the concentration of compound that decreased cell viability by 50% (CC_50_). As a measure of antiviral potential, the selective index (SI) values were calculated as the CC_50_/EC_50_.

A549 cells were plated at 8 x 10^3^ cells per well and HFF cells were plated at 10^4^ on 96-well plates 24 h pre-infection. On the day of infection, USC-093, USC-093D, and cidofovir (CDV) were serially diluted, and aliquots were transferred onto the plates with the cells. The final row on the plates was a no drug control row. Virus at the appropriate concentration for each serotype and cell line was added immediately after drug additions; four replicates were used for each drug concentration. HAdV-infected plates were fixed with 3.7% paraformaldehyde in PBS at 48 h p.i. and stained for the presence of the adenoviral hexon protein using an antibody (mouse monoclonal 2-Hx-2) recognizing hexon from all mammalian adenoviruses, and then visualized with KPL TrueBlue Peroxidase Substrate (SeraCare Life Sciences, Inc.). The assay was read quantitatively on an EliSpot CTL plate reader and graphed in GraphPad Prism 9.

### 2.5 Drug formulation

For in vivo experiments, USC-093 or USC-093D was dispersed in 0.5% aqueous low viscosity carboxymethylcellulose sodium salt, (CMC; Sigma-Aldrich)), then sonicated at room temperature until visual homogeneity (Branson 2510 sonicator) [28]. No loss of homogeneity could be observed visually after 14 days.

### 2.6 Challenge of hamsters with adenovirus; treatment with antiviral compounds

All studies were performed following federal and institutional regulations and were approved by the Institutional Animal Care and Use Committee of Saint Louis University. Male Syrian hamsters with approximate 100 g body weight (*Mesocricetus auratus*) were purchased from Envigo (Indianapolis, IN). Male animals were used because these are more susceptible to human adenovirus infection compared to females [30].

#### 2.6.1 Determining the Maximum Tolerated Dose of USC-093D in Immunosuppressed Syrian Hamsters

Approximately 100 g male hamsters were purchased from Envigo. All hamsters were immunosuppressed using cyclophosphamide (CP). CP was administered intraperitoneally at a dose of 140 mg/kg, and then twice weekly at a dose of 100 mg/kg. USC-093D has shown anti-adenoviral efficacy in vitro. In preparation to test the in vivo efficacy of the compound, we determined the maximum tolerated dose (MTD) of USC-093D and compared it to that of a similar compound, USC-093. Based on existing data, we used dose levels of 10, 20, and 40 mg/kg orally (p.o.), and administered the compound daily for 13 days. There was a total of 7 groups of animals, 6 hamsters each. Hamsters were dosed with vehicle, USC-093D, or USC-093 (**Table 1**). Hamsters were observed and weighed daily. The animals were sacrificed on day 13 and gross pathologic observations were made. Serum samples were collected for analysis of transaminase, creatinine, and blood urea nitrogen levels.

**Table 1.**
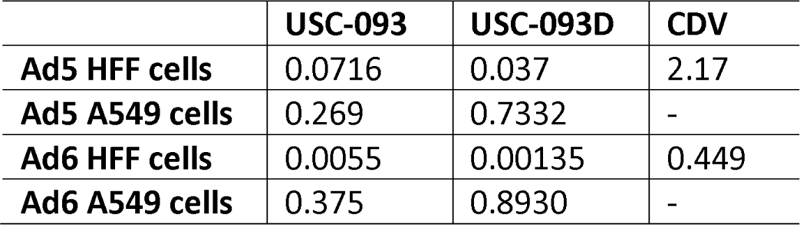
In vitro EC_50_ values (μM).

|  | **USC-093** | **USC-093D** | **CDV** |
| --- | --- | --- | --- |
| **Ad5 HFF cells** | 0.0716 | 0.037 | 2.17 |
| **Ad5 A549 cells** | 0.269 | 0.7332 | - |
| **Ad6 HFF cells** | 0.0055 | 0.00135 | 0.449 |
| **Ad6 A549 cells** | 0.375 | 0.8930 | - |

#### 2.6.2 Determining the Prophylactic Efficacy of compounds USC-093 and USC-093D Against Intranasally Administered Human Type 6 Adenovirus in Immunosuppressed Male Syrian hamsters

Detailed methods were previously explained by Tollefson et al. [23]. Briefly, all hamsters were immunosuppressed using cyclophosphamide (CP). CP was administered intraperitoneally at a dose of 140 mg/kg, and then twice weekly at a dose of 100 mg/kg. USC-093 and USC-093D were obtained in powdered form, suspended in 0.5% carboxymethyl cellulose (Sigma C5678) at the appropriate concentration and sonicated to visual homogeneity. The drugs were made up in a single batch, aliquoted, and stored at 4°C. Approximately 100 g hamsters were dosed with 1 ml volume of suspension. The animals were distributed into 7 groups (**Table 2**). Groups 2 and 3 contained 6 hamsters, while groups 1 and groups 4 to 7 had 15 animals. The hamsters were instilled i.n. with vehicle (Groups 1-3) or 4X10^10^ PFU/kg of HAdV-C6 (Lot# 220719) (Groups 4-7). For all groups, drug administration started 1 day before challenge and continued for the duration of the study. Valganciclovir (VGCV) at 200 mg/kg twice daily (b.i.d.) p.o. was used as positive control. The body weights and any signs of morbidity of the animals were recorded daily. At 3 days post challenge, 5 hamsters (designated at the start of the experiment) from the Vehicle-Vehicle group and all the HAdV-C6-infected groups were sacrificed, and gross pathological observation was performed. Lung samples were collected, and the virus burden was determined by TCID_50_ assay. The remaining 10 hamsters were sacrificed at 7 days post challenge.

**Table 2.** Determining the MTD of USC-093D in immunosuppressed hamsters.

| Groups |  | Drug dose | n | Work Performed |  |
| --- | --- | --- | --- | --- | --- |
|  |  |  |  | At sacrifice | Daily |
| 1 | Drug vehicle | none | 6 | Gross pathology, collect serum for optional transaminase creatinine, and BUN levels | Observation, Body Weight |
| 2 | USC-093D | 10 mg/kg q.d. | 6 |  |  |
| 3 |  | 20 mg/kg q.d. | 6 |  |  |
| 4 |  | 40 mg/kg q.d. | 6 |  |  |
| 5 | USC-093 | 10 mg/kg q.d. | 6 |  |  |
| 6 |  | 20 mg/kg q.d. | 6 |  |  |
| 7 |  | 40 mg/kg q.d. | 6 |  |  |

**Table 3.** Prophylactic study for USC-093 and USC-093D in HAdV-C6-infected, immunosuppressed Syrian hamsters.

| Group # | HAdV-C6 | Compound | Dose | n | Work Performed |  |  |
| --- | --- | --- | --- | --- | --- | --- | --- |
|  |  |  |  |  | Daily | At Necropsy |  |
|  |  |  |  |  |  | Day 3 | Day 7 |
| 1 | – | none | N/A | 5+10 | Dose<br>drug/drug<br>vehicle daily,<br>Observation,<br>Body Weight |  | Sacrifice all |
| 2 | – | USC-093 | 20 mg/kg<br>p.o. q.d. | 6 |  |  |  |
| 3 | – | USC-093D | 20 mg/kg<br>p.o. q.d. | 6 |  |  |  |
| 4 | + | none | N/A | 5+10 |  | lung TCID50 |  |
| 5 | + | USC-093 | 20 mg/kg<br>p.o. q.d. | 5+10 |  |  |  |
| 6 | + | USC-093D | 20 mg/kg<br>p.o. q.d. | 5+10 |  |  |  |
| 7 | + | VGCV | 200 mg/kg<br>b.i.d. | 5+10 |  |  |  |

### 2.7 Statistical analysis

Statistical analysis was performed using GraphPad Prism 4 (GraphPad Software). Two-way ANOVA was used to compare body weight changes. For serum transaminase levels and virus burden in the liver and lung, the overall effect was calculated using Kruskal-Wallis test, and comparison between groups was performed using Mann-Whitney U-test. P ≤ 0.05 was considered significant.

## 3. RESULTS

### 3.1 Synthesis of USC-093/USC-093D

The side chain of homoserine is long and flexible enough to allow the primary hydroxyl group to attack the carbonyl. Instead of coupling with the lipophilic amine, Boc-protected homoserine undergoes intramolecular cyclization. Therefore, the synthesis of USC-093 began with commercially available

-L-homoserine lactone (**Scheme 1**). Treatment of Boc-L-homoserine lactone 1 with hexadecylamine **2** in THF at 60 °C over two days, which leads to the opening of γ-lactone ring yielding the amide **3** in good yield.

The amide **3** was combined with (*S*)-HPMPA **4** in DMF, followed by the addition of DIEA and PyBOP. The PyBOP mediated coupling reaction was carried out at 40°C for 2 h, which furnished oxaphosphinanes **5**. In theory, two equivalents of PyBOP would be essential for intermediate formation. The first equivalent of PyBOP is used to convert HPMPA into cyclic HPMPA. Then, the second equivalent reacts with the cyclic HPMPA to produce an HOBt-HPMPA adduct, which is attacked by the hydroxyl on the promoiety to form the oxaphosphinane. However, an excess of PyBOP may be required to compensate for the portion that becomes hydrolyzed during the reaction.

Retention of the cycle to eliminate one of the hydroxyl charges on the phosphonate at physiological pH may seem advantageous; however, cyclic prodrugs have been shown to have several disadvantages compared to their acyclic counterparts. First, cyclic prodrugs create a stereocenter on the phosphorous (Sp and Rp) creating two different diastereomers. These diastereomers have different stabilities, conformations, and half-lives [31, 32]. Purifying diastereomers is difficult and time consuming whereas using a hydrolysis reaction eliminates the stereocenter efficiently. The cyclic prodrugs are also disadvantaged due to their shorter half-life when compared to acyclic prodrugs. The brevity of their half-life may cause the prodrug to degrade before crossing the cellular membrane [31, 32]. We therefore chose to generate the acyclic intermediate by dissolving the oxaphosphinane **5** in MeCN, then treating the solution with NH_4_OH in excess and heating to 45°C over six days, giving the penultimate phosphonate **6** in moderate yield.

In the final N-terminal amino deprotection step, trifluoroacetic acid (TFA) was initially used to remove the Boc protecting group, yielding the desired prodrug in its TFA salt form. To ensure safety for subsequent in vitro and in vivo studies, it was necessary to convert the TFA salt to a less toxic form [33]. We therefore treated the prodrug with concentrated NH_4_OH followed by purification via silica column chromatography. However, this process had to be repeated multiple times, and it was not always possible to remove the TFA. Consequently, we changed to concentrated HCl for deprotection of the amino group, resulting in formation of the prodrug as an HCl salt, a safer and widely accepted formulation encountered in many FDA-approved drugs [34] [35]. The final product (USC-093 **7**) was then obtained in good yield by stirring phosphonate **6** in THF with saturated aq. HCl at room temperature overnight.

USC-093D was prepared by the same route starting from Boc-D-homoserine lactone.

### 3.6 In vitro studies comparing USC-093 and USC-093D

With USC-093’s promising results from our previous study [23], further SAR studies were performed by comparing USC-093 and its D-isomer, USC-093D. Using the UAB assay, the D-isomer exhibited an EC_50_ of 0.03 µM and CC_50_ of > 0.40 µM with an overall SI_50_ of 22 against HAdV-C5 in HFF cells, and in comparison, the L-isomer, USC-093, displayed an EC_50_ of 0.05 µM and CC_50_ of > 0.40 µM with an overall SI_50_ of 23. Concomitant assays of the control agent CDV demonstrated an EC_50_ of 7.94 µM and CC_50_ of 117.6 µM. Both isomers exhibited good antiviral potency against HAdV-C5, with the D-isomer having a slightly lower EC_50_ than the L-isomer.

To further evaluate USC-093 and USC-093D, in vitro assays were run at SLU with HAdV-C5 and HAdV-C6 in HFF cells (**Table 1, Figure 5**). Because HAdV-C6 replicates better in Syrian hamsters than HAdV-C5 and therefore is used as the challenge virus in in vivo assays [25, 26], HAdV-C6 is used in in vitro studies of the most promising drug candidates. Both types also were tested in A549 cells. While HFF cells exhibit lower infectivity and viral replication, A549 cells are at the highest end of infectivity and viral replication. Thus, these two cell lines cover the range of permissivity for HAdV. A549 cells typically require more substantial drug concentrations for inhibition of replication (**Table 1, Figure 5**).

### 3.7 USC-093 and USC-093D maximum tolerated dose

At the 10 and 20 mg/kg dose levels, both compounds caused a small but significant delay in weight gain (**Figure 3**). At 40 mg/kg, USC-093 prevented weight gain, while USC-093D caused weight loss. Hamsters in both 40 mg/kg groups had uncharacteristically dilute urine.

**Figure 3.**
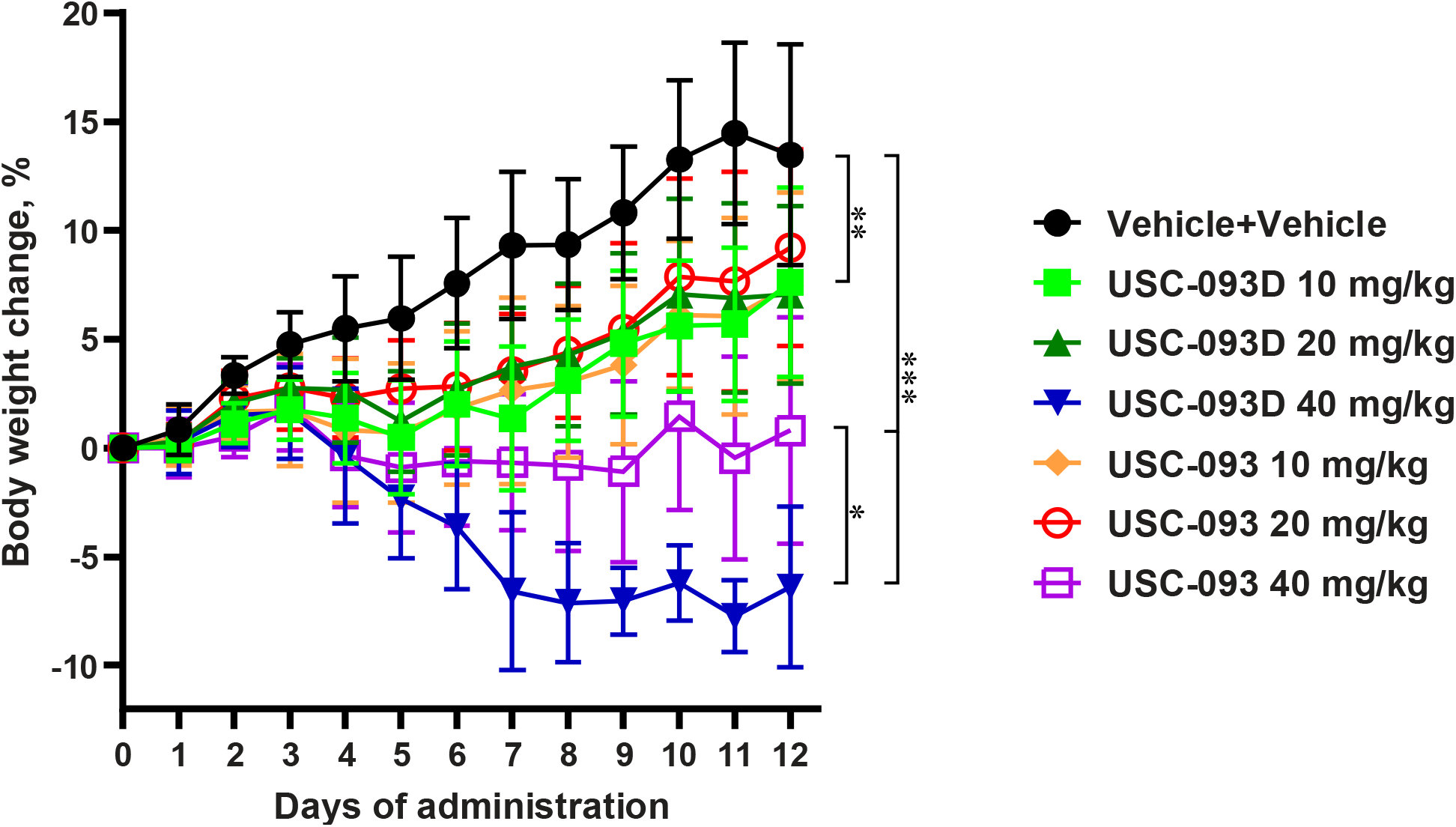
USC-093 and USC-093D at 10 and 20 mg/kg of and USC-093 at 40 mg/kg decreased the growth rate of hamsters, while 40 mg/kg of USC-093D caused weight loss. The symbols represent the group mean, and the error bars signify the standard deviation. ^*^: p<0.05, ^**^: p<0.01, ^***^: p<0.001 (two-way ANOVA).

After 13 days of dosing, the animals were sacrificed and necropsied. Five hamsters in the USC-093D group had moderately pale kidneys, while one hamster had severely pale kidneys. Three of six hamsters in the USC-093 40 mg/kg group had minimally pale kidneys. No significant gross necropsy findings were noted for the groups dosed with 10 or 20 mg/kg of either compound.

Hamsters in the groups that received 40 mg/kg of either drug had decreased blood urea nitrogen and increased chloride levels. The creatinine and liver transaminase levels were normal (**Figure 4, Table 2**).

**Figure 4.**
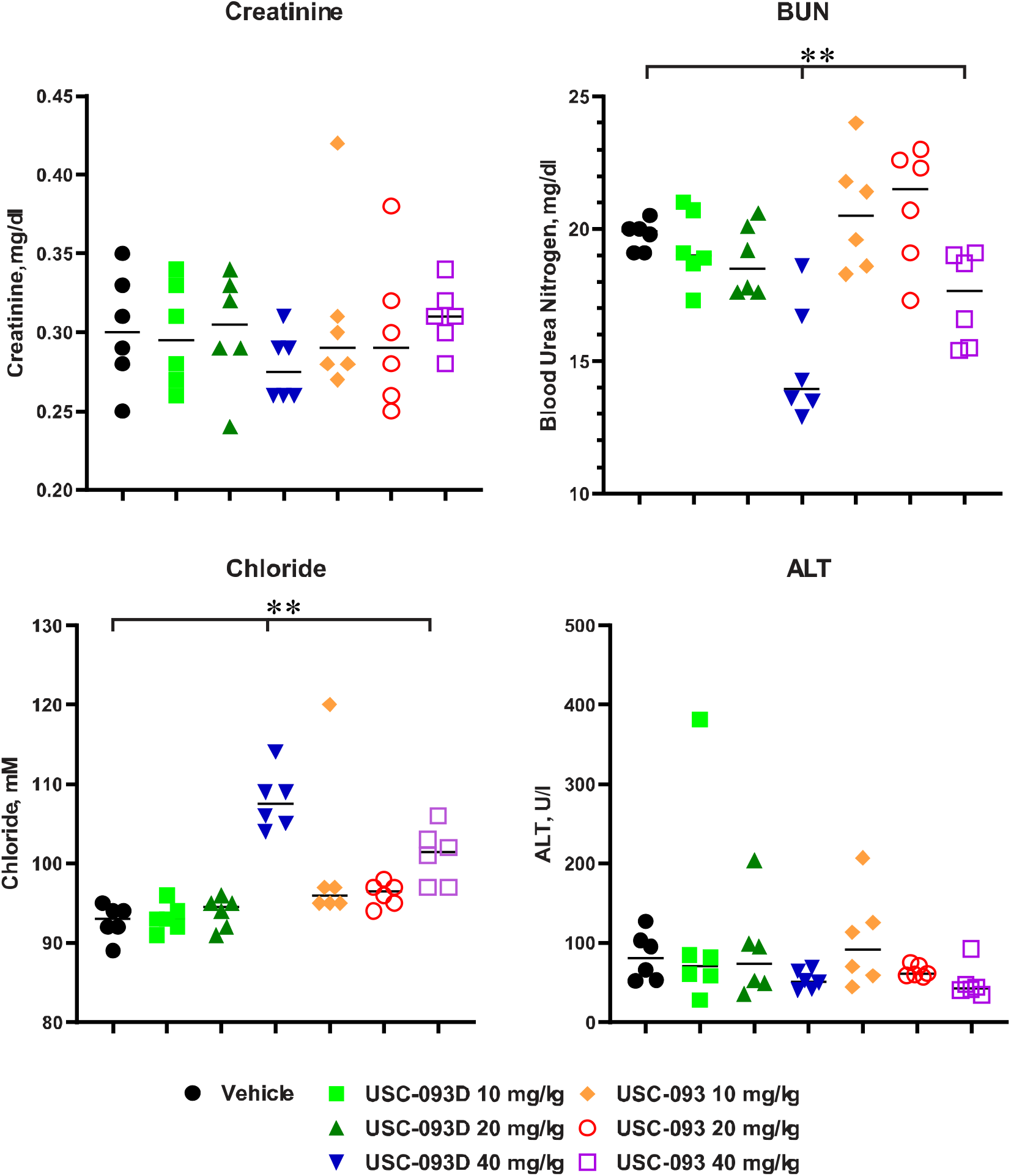
At 40 mg/kg, USC-093 and USC-093D caused a decrease of blood urea nitrogen levels and an increase of chloride levels. Creatinine and alanine aminotransferase levels were normal. The symbols represent values from individual animals, and the horizontal bars signify the median. ^**^: p<0.01 (Mann-Whitney U-test).

Once daily oral administration of 40 mg/kg of either USC-093 or USC-093D for 13 consecutive days was toxic in immunosuppressed Syrian hamsters. The target for toxicity was the kidney. Note that no increased creatinine levels were seen; this is in contrast with previous data obtained with similar compounds. As of now, we do not have an explanation for this. Of the two compounds, USC-093D was more toxic than USC-093. For both compounds, the MTD for oral administration was established at 20 mg/kg q.d.

#### 3.8 USC-093 and USC-093D had marginal effect against respiratory HAdV-C6 infection in vivo

Immunosuppressed hamsters were infected intravenously with HAdV-C6. No treatment-related deaths presented in the study (**Figure 5**). Both USC-093 and USC-093D were dosed at the maximum tolerated dose (MTD), which was 20 mg/kg orally (p.o.) once daily (q.d.). At 3 days post challenge, all hamsters in the HAdV-C6-infected groups had red mottled lung. However, the severity of the lesions was lower in the VGCV-treated control group. At 7 days post challenge, the pathology persisted in the lungs of hamsters in the HAdV-C6+Vehicle and HAdV-C6+USC-093 groups. Three hamsters in both the USC-093D-treated and VGCV-treated groups presented with no lung pathology, and severity of the lesions was lower than that in untreated hamsters in the remaining animals in these groups. As a surrogate measurement for lung pathology (mostly measuring infiltration by immune cells), the lungs of the hamsters sacrificed at 3 days post challenge were weighed and found that lung weights were marginally lower in the USC-093D-treated and VGCV-treated group compared to the HAdV-C6+Vehicle group (**Figure 6**). The virus titers in the lung were measured and found that USC-093 marginally inhibited the virus replication in the lungs of HAdV-C6-infected hamsters at 3 days post challenge; however, the decrease of virus burden was not statistically significant (**Figure 7**).

**Figure 5.**
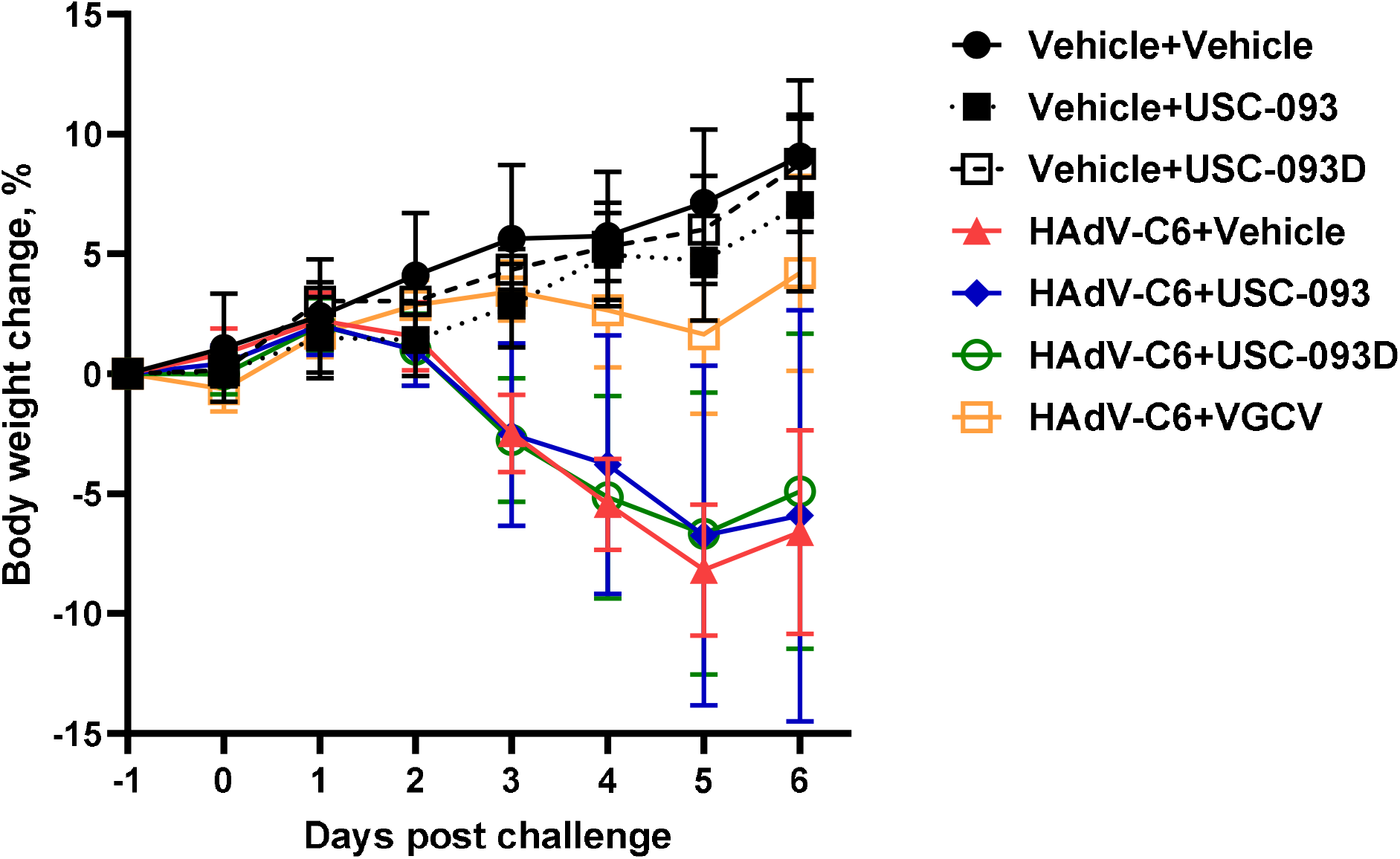
Neither USC-093 nor USC-093D mitigated HAdV-C6-induced morbidity. Each symbol represents the group mean; the error bars depict the standard deviation. HAdV-C6+Vehicle vs. HAdV-C6+VGCV p<0.0001 (two-way ANOVA). Compounds were dosed at 10 mg/kg p.o. q.d. (USC-093 and USC-093D) or 200 mg/kg p.o. b.i.d. (VGCV), starting 1 day before challenge.

**Figure 6.**
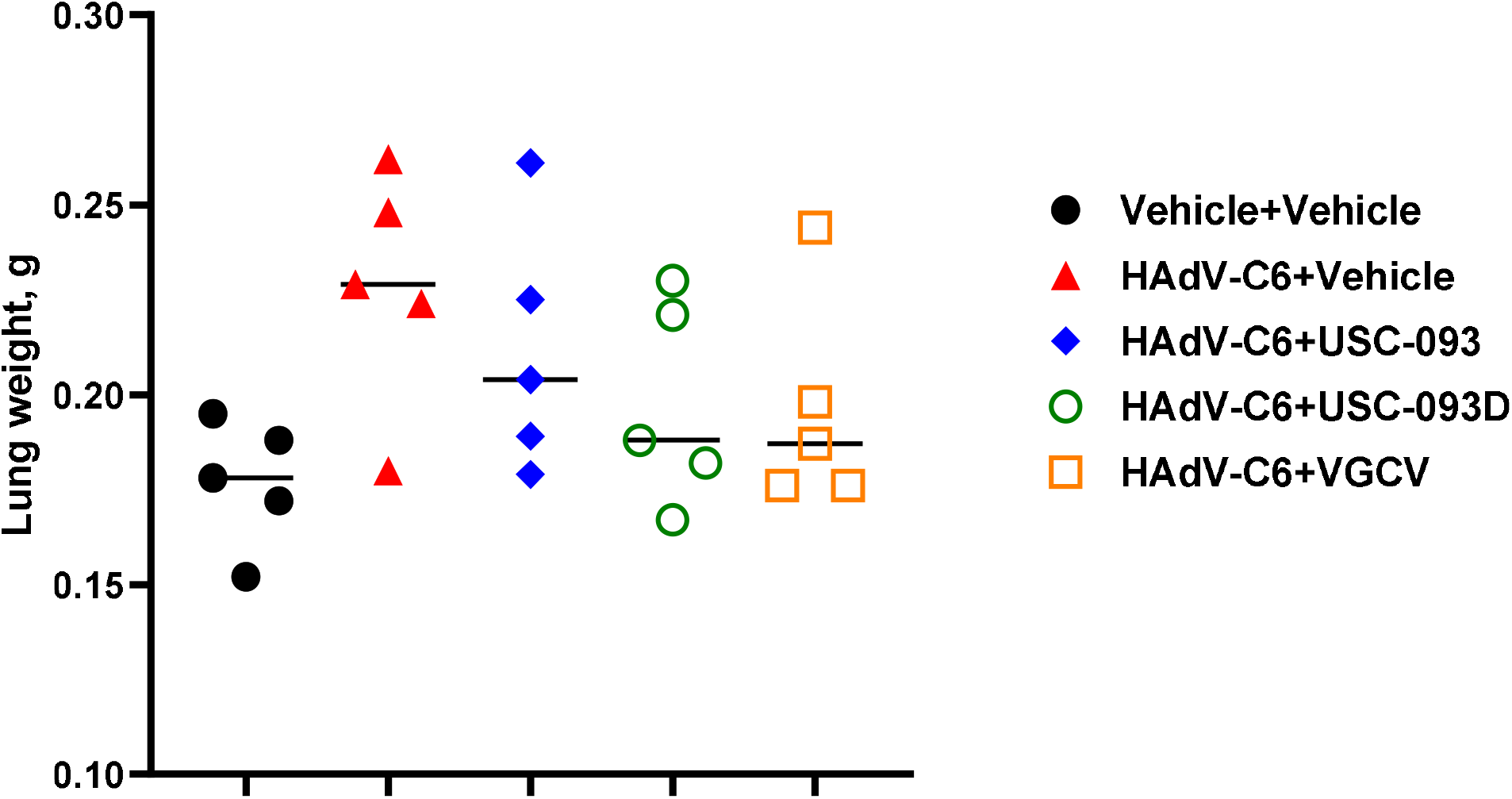
USC-093D marginally mitigates the lung pathology induced by intranasal infection with HAdV-C6. Weight of the left lung lobe is shown; symbols represent data from individual animals, and the bar symbolizes the median. The differences are not statistically significant (Mann-Whitney U-test).

**Figure 7.**
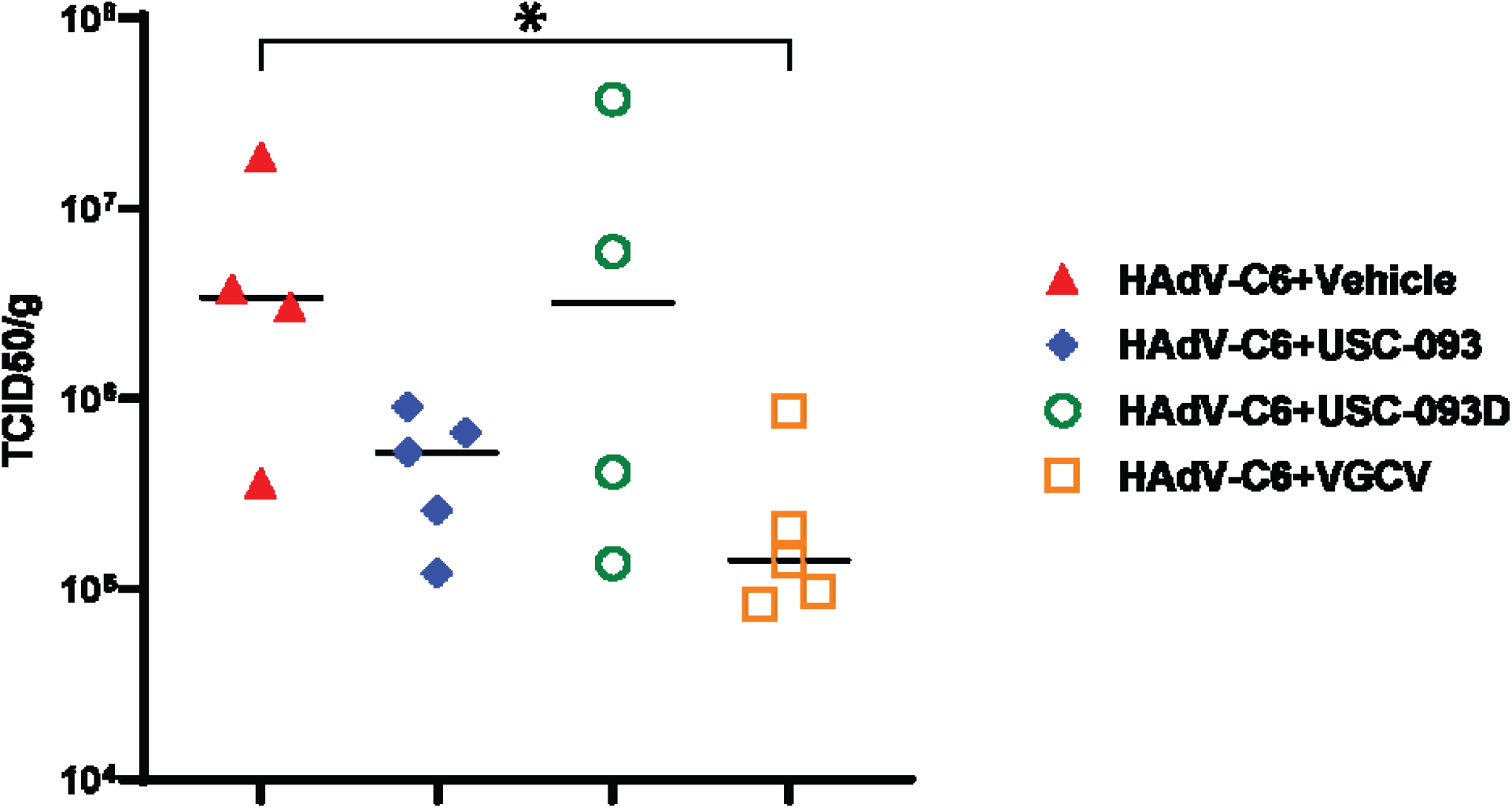
USC-093 marginally inhibits the replication of HAdV-C6 in the lung of immunosuppressed hamsters. The symbols represent data from individual animals, and the bar symbolizes the median. ^*^: p<0.05 (Mann-Whitney U-test).

In conclusion, USC-093D showed comparable results with USC-093. At the 20 mg/kg dose level, the drugs did not significantly reduce lung pathology, but there was a marginal effect by USC-093D. USC-093 inhibited the replication of HAdV-C6 in the lung to some extent; however, the effect was not statistically significant. The in vivo efficacy of the two isomers against HAdV-C6 indicate that the stereochemistry of homoserine has limited effect on drug transportation.

## 4. DISCUSSION

With the increase of patients on drastic immunosuppressive regimes and the high mortality rate caused by viral infections in these patients, the need to develop effective antiviral drugs is obvious. This need is especially urgent in the case of adenoviruses, as presently there is no specific FDA-accepted drug to treat adenovirus infections. Further, emerging adenovirus strains can cause mortality in otherwise healthy humans, and HAdV-F41, an enteric adenovirus, may be implicated in acute pediatric hepatitis cases [3, 36, 37]. Cidofovir, an acyclic nucleoside phosphonate drug is often used off-label to treat patients with multi-organ adenovirus infections; however, the nephrotoxicity of this compound presents a serious drawback for this therapy. We have initiated a project to improve the pharmacology of acyclic nucleotide phosphonate compounds by linking an alkane chain to the parental compound to improve bioavailability.

To compare the in vivo activity of the two compounds, we used the immunosuppressed Syrian hamster model. These animals are permissive for infections with species C human adenoviruses and develop pathology similar to human patients [38]. Species C human adenoviruses are often isolated from immunosuppressed patients, and thus are a relevant challenge strain [11].

In the development of prodrugs, significant research has been conducted on the use of both D- and L-amino acids to optimize drug delivery and efficacy [39]. The use of D-amino acids has been highlighted in gemcitabine prodrug design, where the D-conformation amino acids have been shown to confer better permeability and greater stability in cell homogenates, slowing enzymatic bioconversion rates which can be beneficial for maintaining therapeutic drug levels over extended periods [40]. Conversely, some studies also evidenced that L-amino acids are preferred for high permeability, for example, L-valyl ester prodrug of acyclovir (L-Val-ACV) has ten-fold higher permeability than its D-isomer [41]. Moreover, L-amino acids have demonstrated superior results in enhancing bioavailability and pharmacokinetic profiles, as seen in drugs like gemcitabine, the 5’-L-valyl and 5’-L-isoleucyl monoester prodrugs exhibited increased uptake in HeLa/hPEPT1 cells compared to HeLa cells and chemical stability in buffers [42]. The controversial conclusions for L- and D-amino acids in prodrug strategy indicates no considerable preference for one over the other. Given these mixed results, it is crucial to conduct a thorough SAR study investigating both L- and D-amino acids in our prodrug to assess their impact on drug delivery and efficacy. However, our data demonstrate that USC-093 and USC-093D, which differ only in their stereochemistry, exhibit similar potency in vivo. This suggests that the stereochemistry of the amino acid has a minor effect in this case, implying that our prodrug is not actively transported by hPEPT1, as stereoselectivity often plays a key role in transporter-mediated uptake [43]. Thus, the stereochemistry of homoserine in our compounds may not be a critical determinant of efficacy.

## 5. CONCLUSION

As allogeneic hematopoietic stem transplantation continues to be the state of the art for many pediatric hematologic malignancies, the number of severely immunocompromised patients is expected to remain high. Thus, it is imperative that effective anti-adenoviral agents are developed. However, achieving good efficacy is just one aspect of the development. In many cases, promising new compounds failed during clinical trials or even earlier in development because of unacceptable toxicity. Therefore, we place equal emphasis on antiviral efficacy, bioavailability, and good toxicology profile. In conclusion, this study described a detailed synthesis for the ANP prodrugs USC-093 and USC-093D, which differ only in the stereochemistry of their homoserine moiety. Both prodrugs demonstrated potent antiviral activity in vitro against HAdV-C5, with similar potency, and negligible cytotoxicity. In vivo results against respiratory HAdV-C6 infection in immunosuppressed hamsters revealed that the two isomers exhibited comparable efficacy. These findings suggest that the stereochemistry of homoserine has a minor impact on the efficacy of the prodrugs, and their mechanism of transport is likely not dependent on stereoselective pathways, such as hPEPT1-mediated uptake. Further research will be required to optimize the pharmacokinetic and pharmacodynamic profiles of these prodrugs for improved antiviral efficacy in vivo. Such studies will be facilitated by the eminent tunability of our prodrug platform.

## ASSOCIATED CONTENT

### Supporting Information

Spectra including ^1^H NMR, ^31^P NMR, MS of USC-093 and USC-093D. ISCO chromatograms of intermediates, USC-093 and USC-093D.

## AUTHOR INFORMATION

### Co-corresponding Author

^*^

### Present addresses

^a^Saint Louis University School of Medicine, St. Louis, MO 63104, USA

^b^University of Southern California, Los Angeles, CA 90089, USA

### CRediT authorship contribution statement

JY: formal analysis, Investigation, methodology, validation, writing, review and editing

SR: investigation, methodology

JL: investigation, methodology

SF: Investigation, Methodology

YB: writing, review and editing, validation

NL: writing, review and editing

IK: project administration, review and editing

BAK: supervision, methodology, review

CBH: methodology, investigation, writing, review and editing

SHJ: funding acquisition, methodology, supervision, investigation, review and editing,

AET: methodology, investigation, review and editing)

AS-C: investigation

KT: funding acquisition, methodology, investigation, supervision, writing, review and editing

SHJ: funding acquisition, methodology, investigation, review and editing, supervision,

CEM: funding acquisition, project administration, resources, supervision, methodology, review and editing.

## NOTES

The authors declare no competing financial interest.

## ACKNOWLEDGMENTS

We wish to acknowledge Dr. William Wold† and Dr. Mark Prichard† for contributions at the inception of this project.

## ABBREVIATIONS

(HAdV): Human adenoviruses
(CDV): (*S*)-9-(3-hydroxy-2-phosphonylmethoxypropyl)adenine ((*S*)-HPMPA); cidofovir
(BCV): Brincidofovir
(LR-MS): Low-resolution mass spectrometry
(HPLC): high-performance liquid chromatography
(THF): tetrahydrofuran
(EtOAc): ethyl acetate
(PyBOP): benzotriazol-1-yloxytripyrrolidinophosphonium hexafluorophosphate
(DMF): N,N-dimethylformamide
(DIEA): N,N-diisopropylethylamine
(DCM): dichloromethane
(MeOH): methanol
(NH_4_OH): ammonium hydroxide
(MeCN): acetonitrile
(NIAID): National Institute of Allergy and Infectious Diseases
(HFF): human foreskin fibroblast
(CPE): cytopathic effect
(CMC): carboxymethylcellulose sodium salt,
(CP): cyclophosphamide
(i.v.): intravenously
(p.o.): (PFU)/kg (approximately the 50% lethal dose); oral gavage
(VGCV): Valganciclovir
(b.i.d.): twice daily
(TCID_50_): 50% Tissue Culture Infectious Dose .

## FIGURES, SCHEMES AND CAPTIONS

**Scheme 1.**
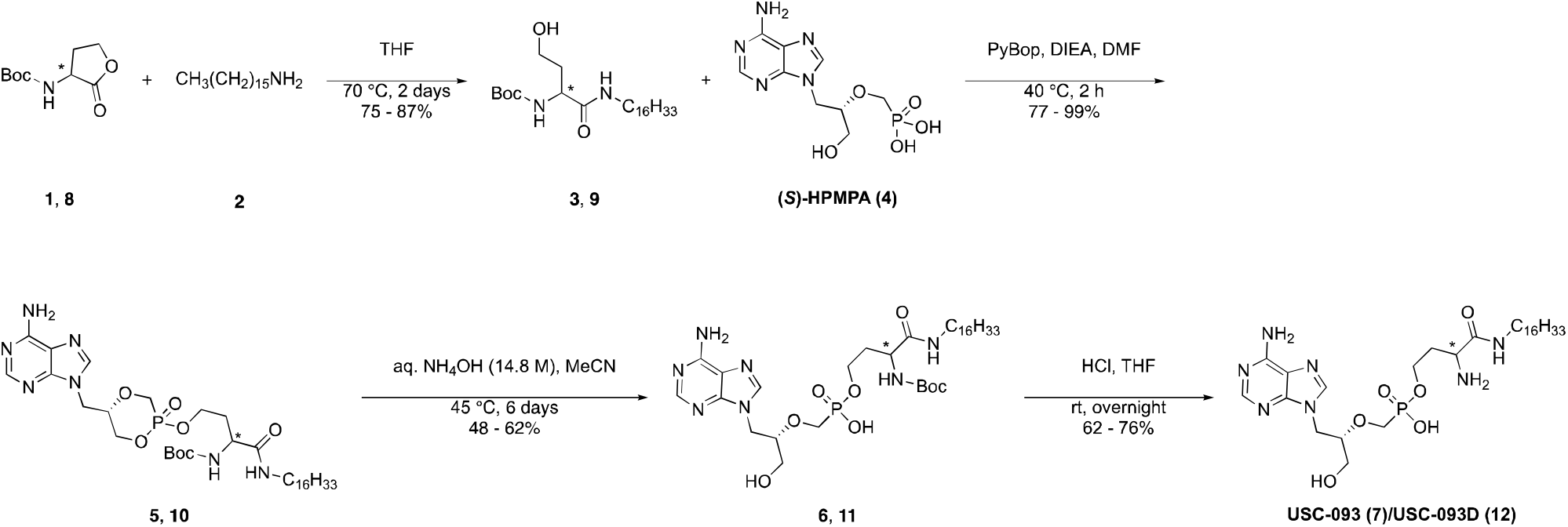
Chemical synthesis of USC-093 and USC-093D.

