## Supplementary Information for "Synthesis of USC-093 and comparison with its promoiety enantiomer USC-093D against adenovirus in vitro and in a Syrian hamster model"

Table S1. Serum Chemistry.……………………………………………….………………………………………..……………………....28

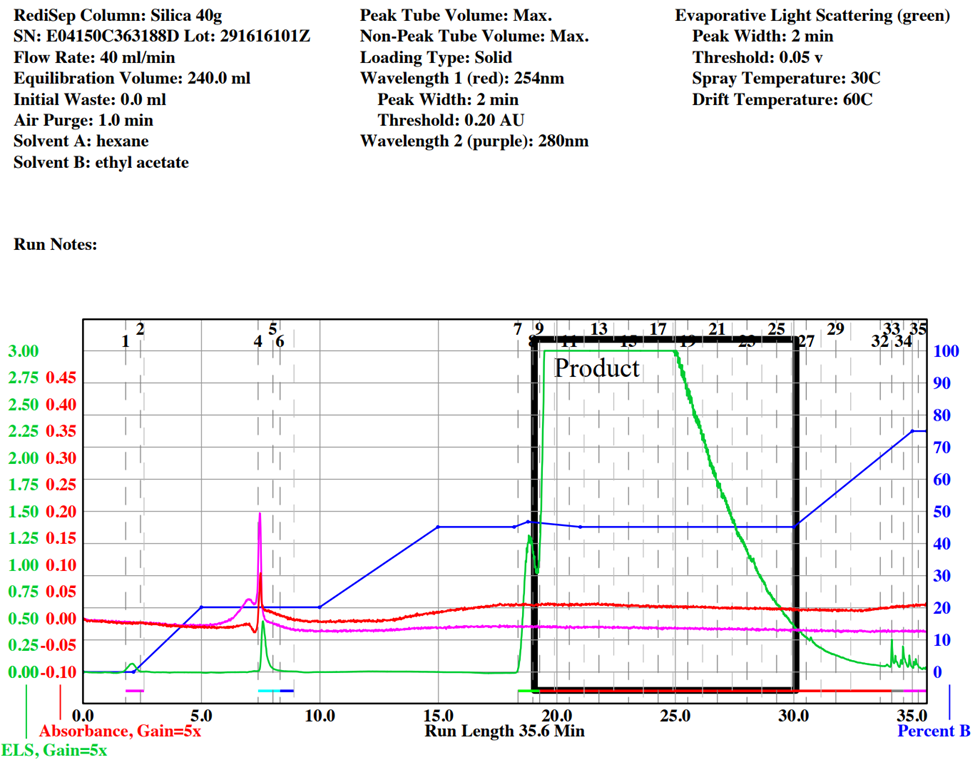

Figure S1. ISCO chromatogram for the promoiety (**3**).

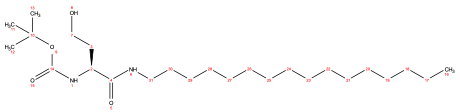

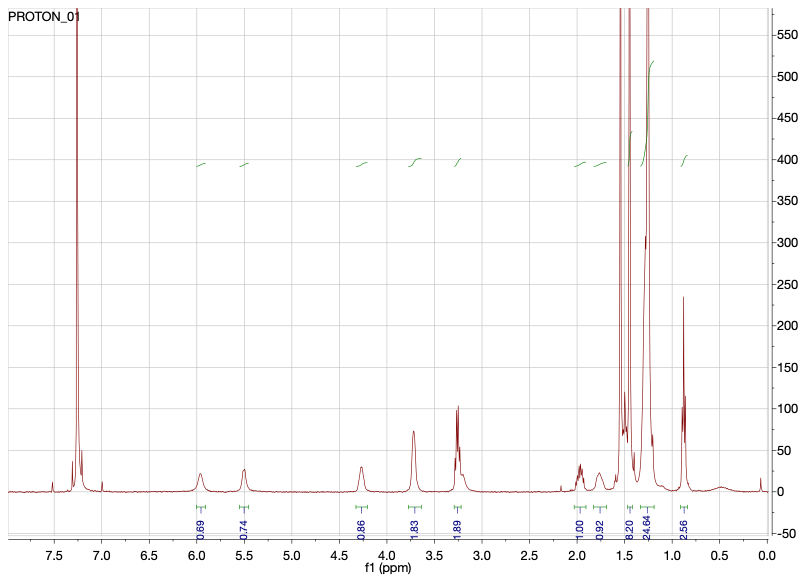

Figure S2. ^1^H NMR (400 MHz, CDCl_3_) spectrum of the promoiety (**3**).

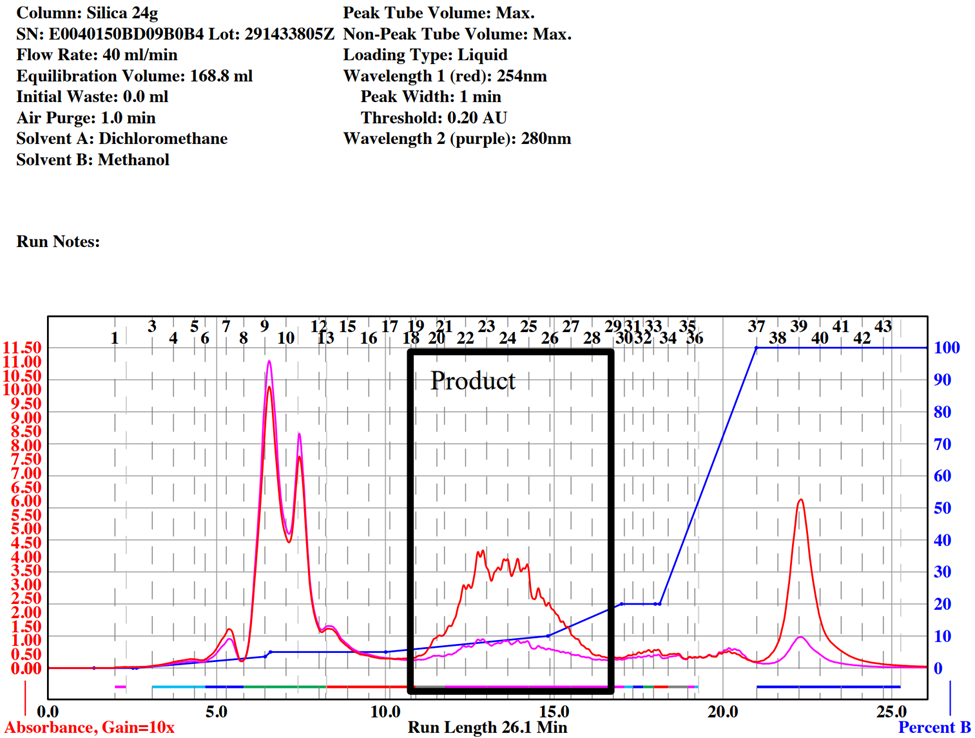

Figure S3. ISCO chromatogram for oxaphosphinanes (**5**).

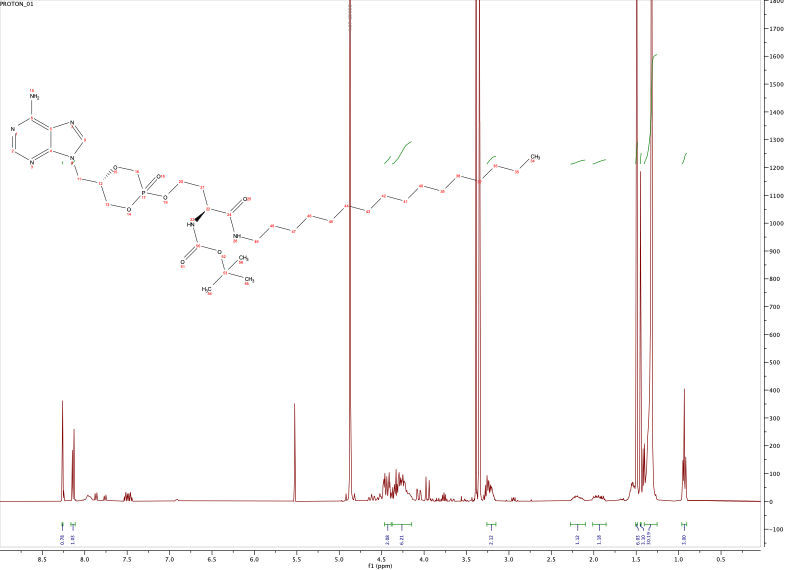

Figure S4. ^1^H NMR (400 MHz, CD_3_OD) spectrum of oxaphosphinanes (**5**).

Trace peaks in the aromatic region are from side products which was removed via flash chromatography in the next step.

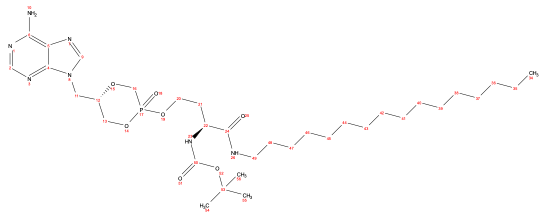

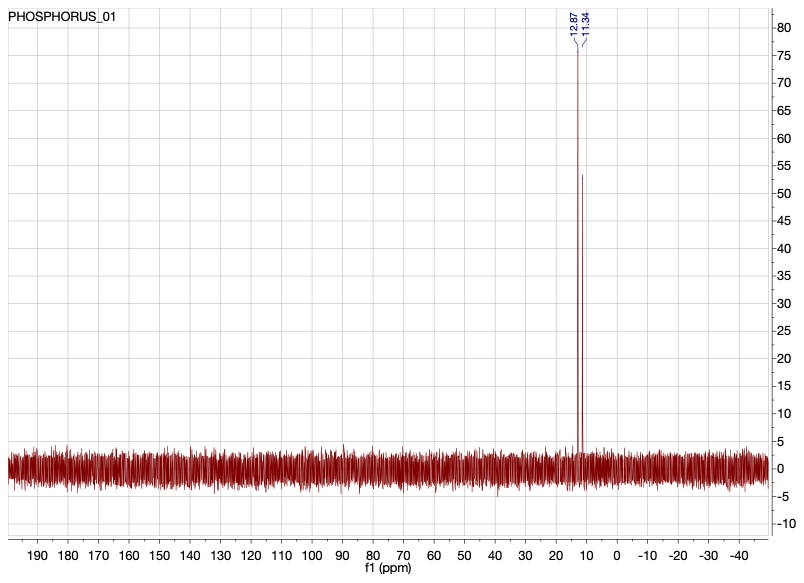

Figure S5. ^31^P NMR (162 MHz, CDCl_3_) spectrum of oxaphosphinanes (**5**).

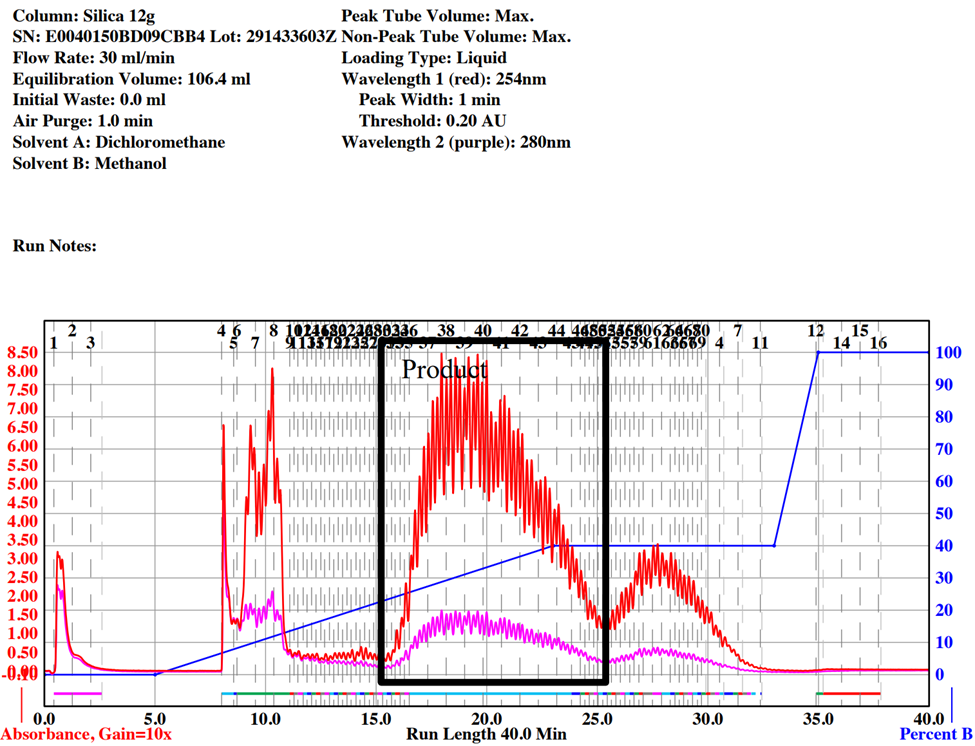

Figure S6. ISCO chromatogram for phosphonate (**6**).

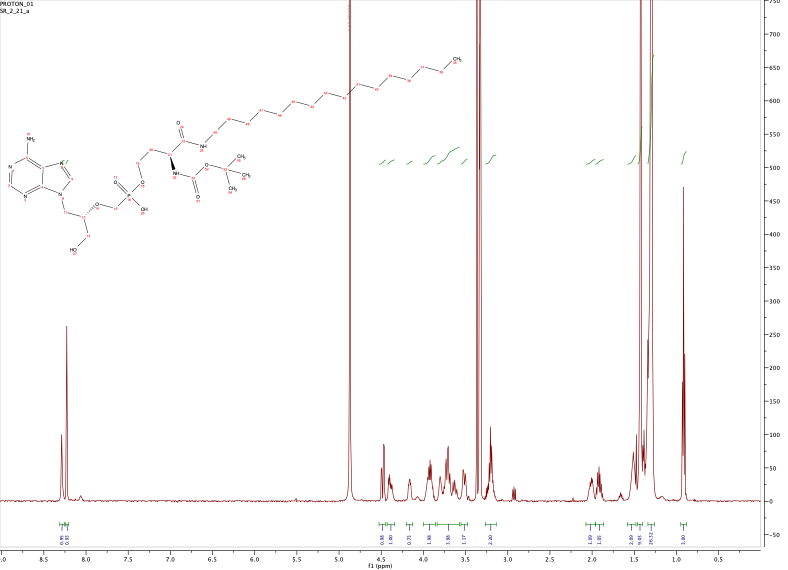

Figure S7. ^1^H NMR (500 MHz, CD_3_OD) spectrum of phosphonate (**6**).

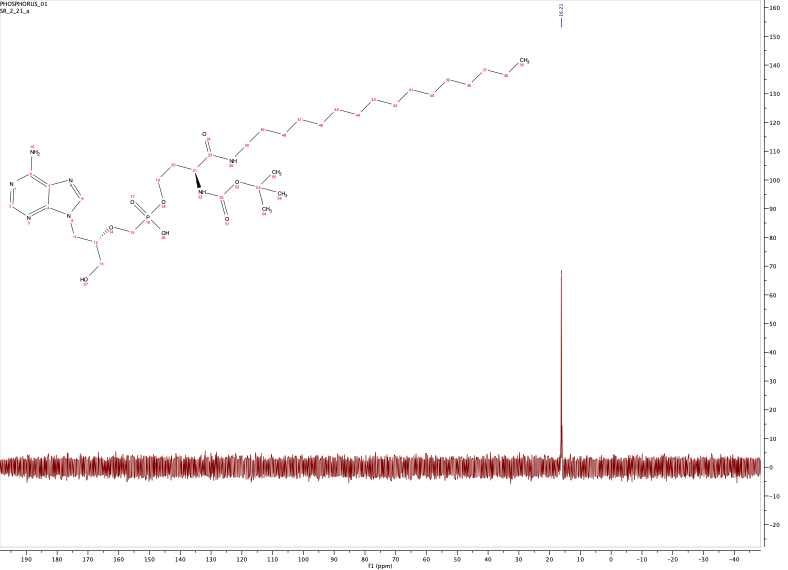

Figure S8. ^31^P NMR (202 MHz, CD_3_OD) spectrum of phosphonate (**6**).

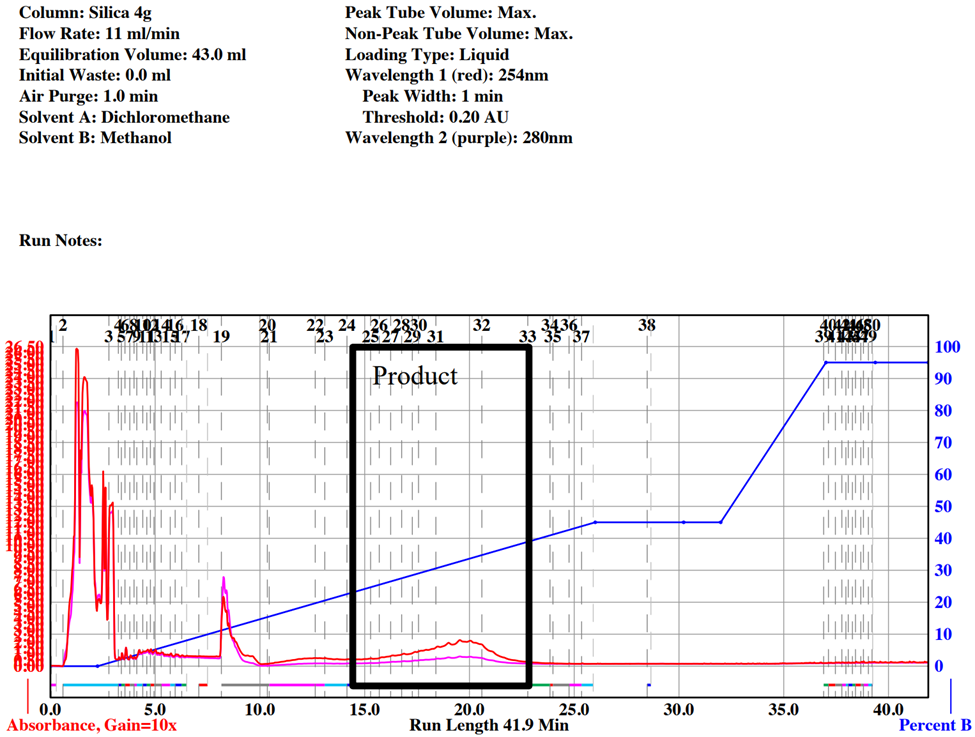

Figure S9. ISCO chromatogram for USC-093 (**7**).

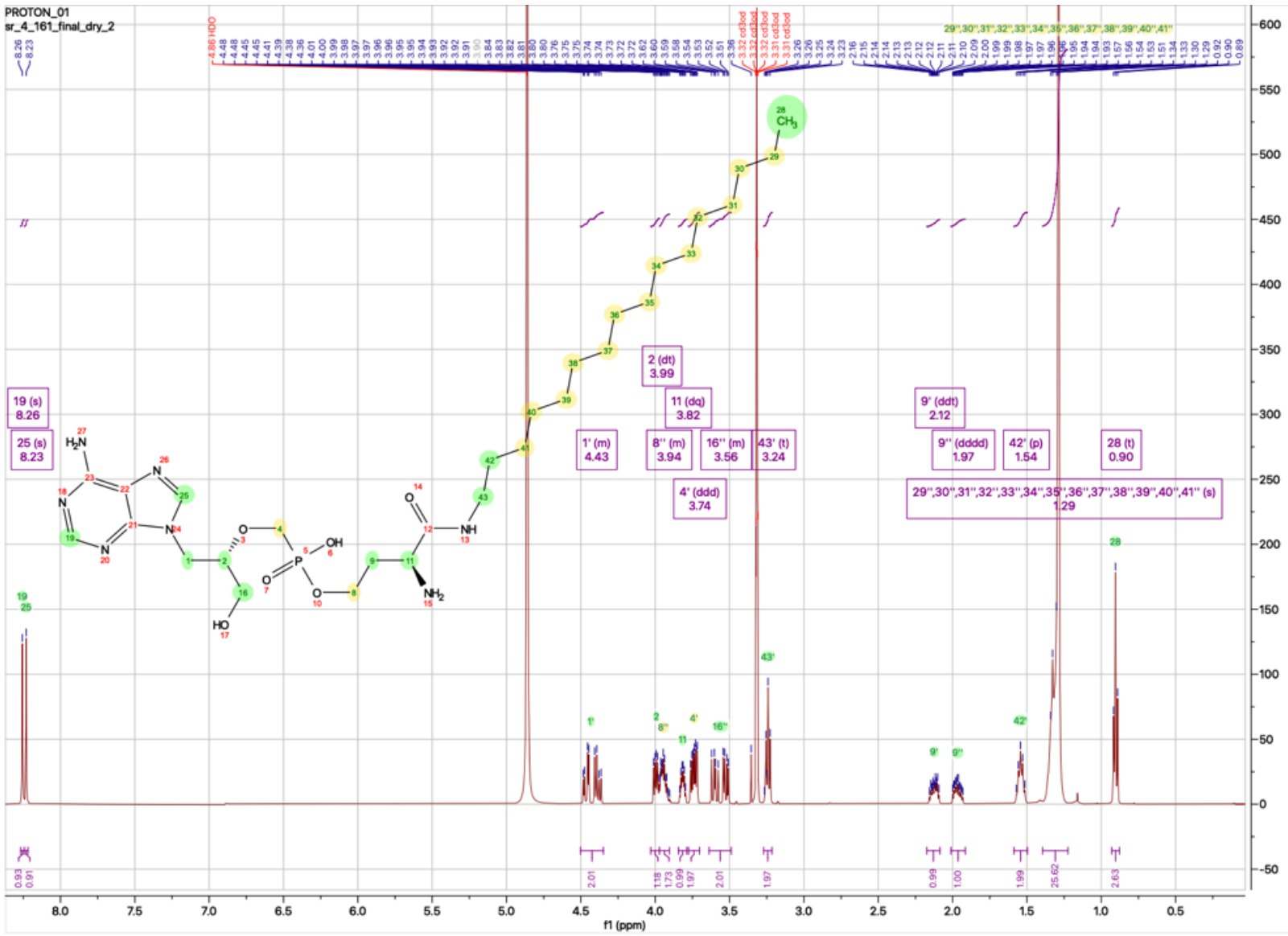

Figure S10. ^1^H NMR (500 MHz, CD_3_OD) spectrum of USC-093 (**7***)*.

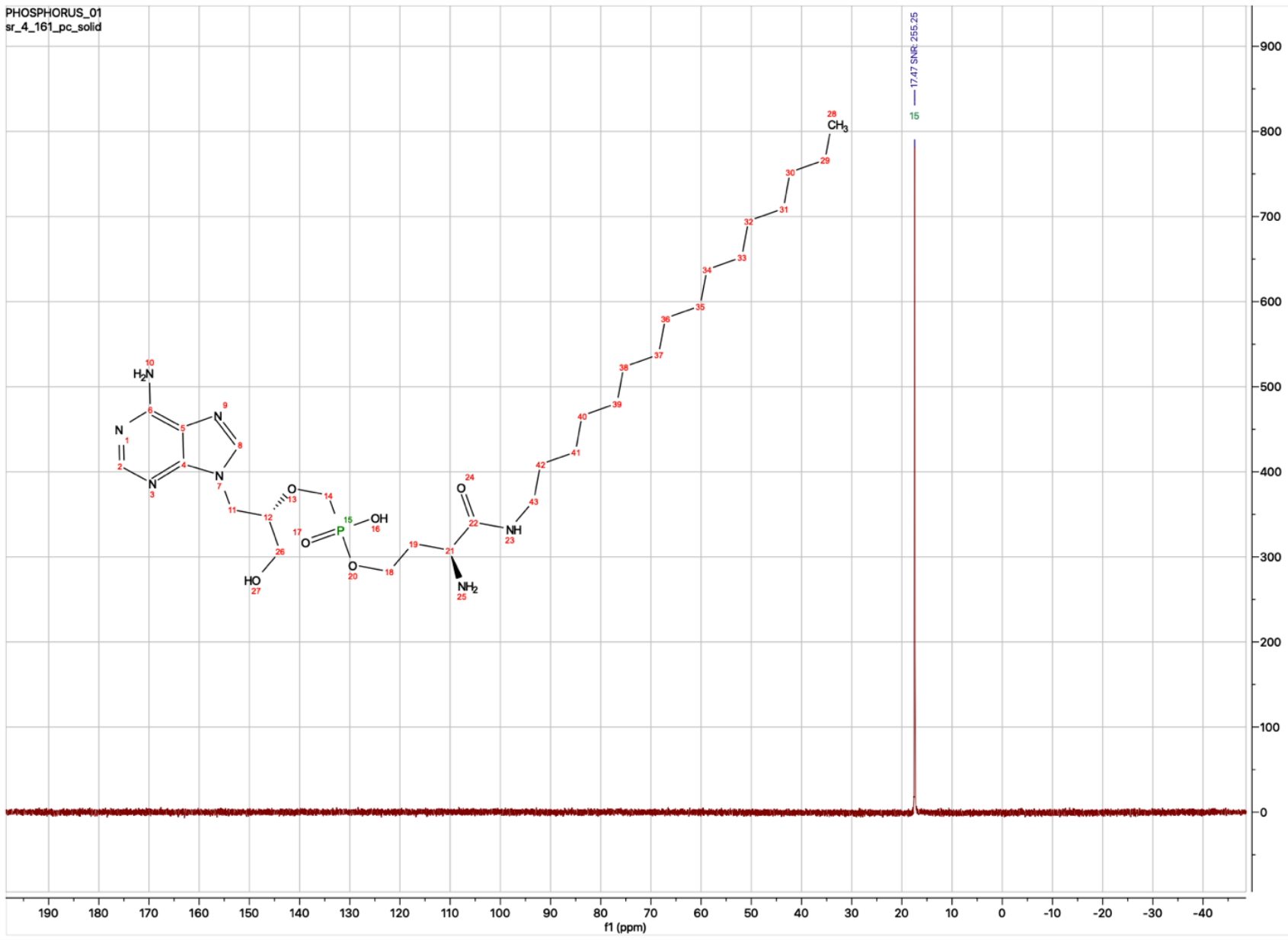

Figure S11. ^31^P NMR (162 MHz, CD_3_OD) spectrum of USC-093 (**7**).

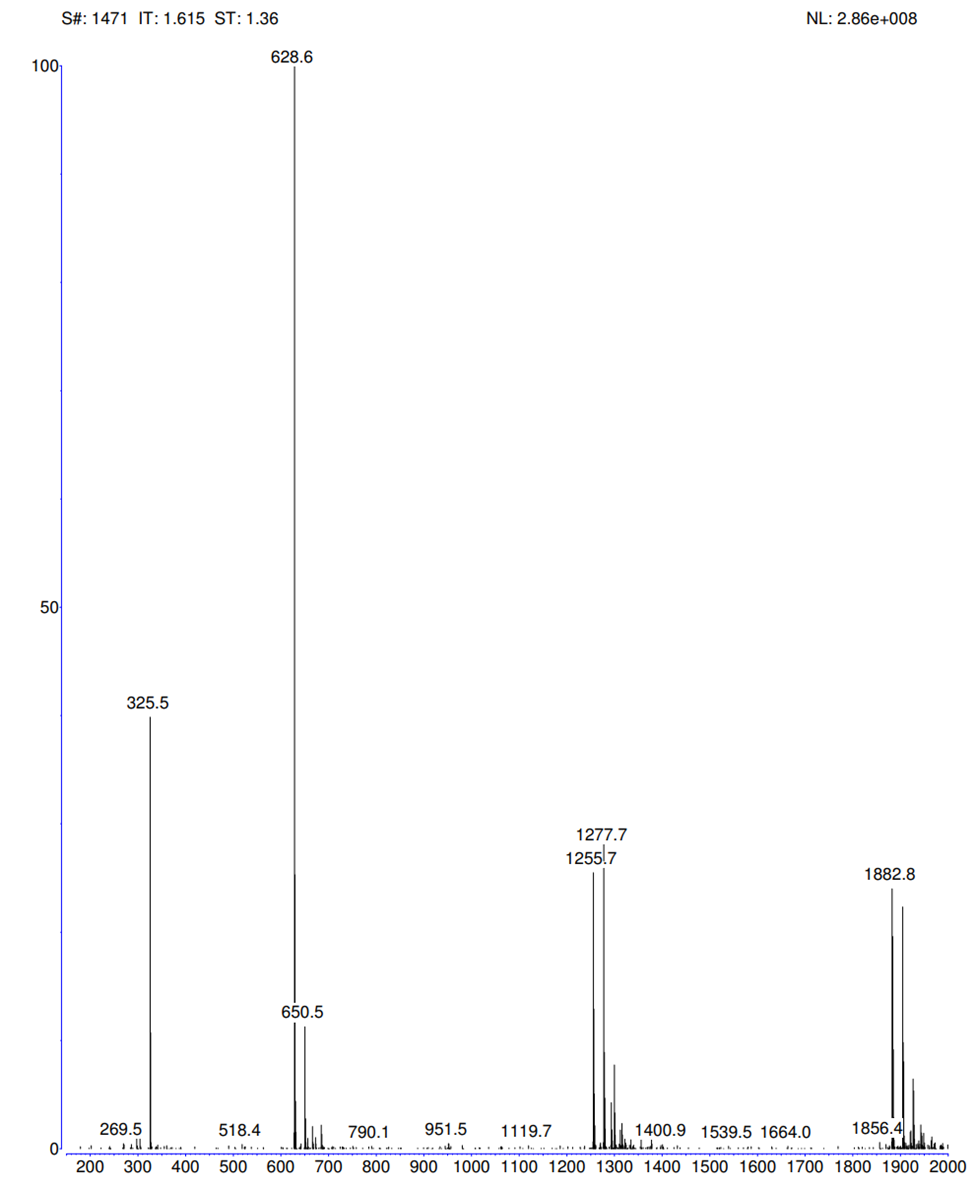

Figure S12. MS spectrum of USC-093 (**7**) in positive mode.

[M+H]^+^: C_29_H_55_N_7_O_6_P ^+^; Calcd: 628.39; found: 628.6 – Thermo-Finnigan

| Analysis | Result | Calculated |
| --- | --- | --- |
| C: Carbon | 55.19 | 55.49 |
| H: Hydrogen | 8.73 | 8.67 |
| N: Nitrogen | 15.08 | 15.62 |

Figure S13. CHN analysis of USC-093 (**7**).

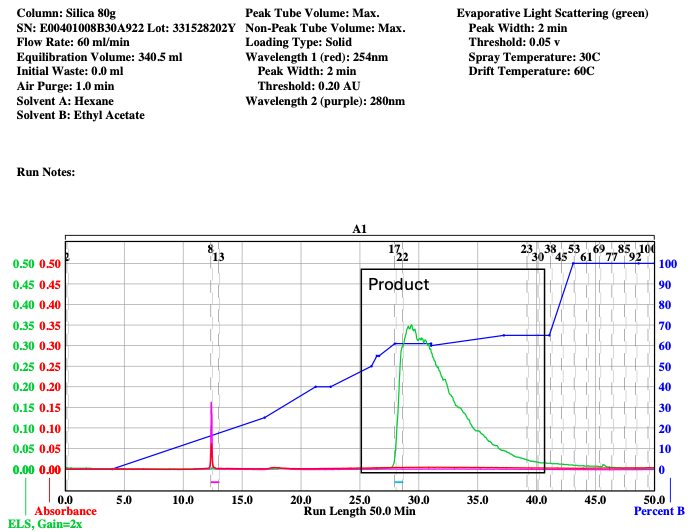

Figure S14. ISCO chromatogram for promoiety (**9**).

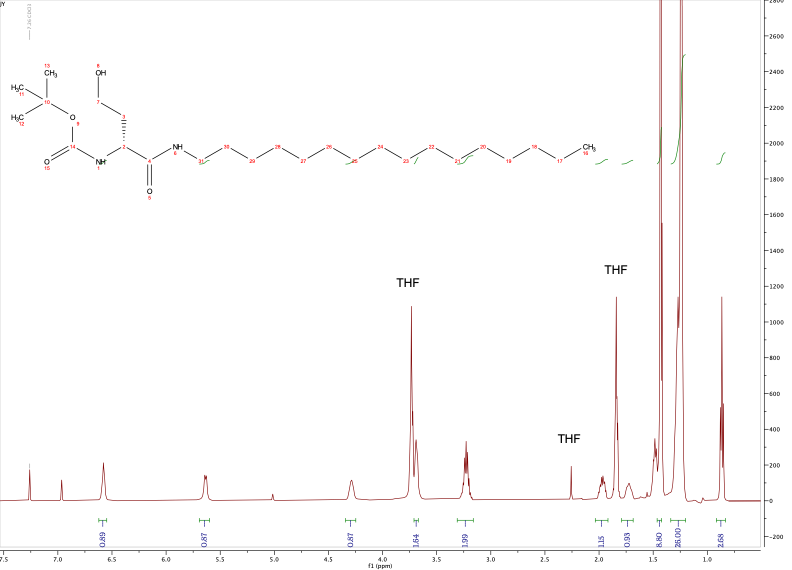

Figure S15. ^1^H NMR (500 MHz, CDCl_3_) spectrum of promoiety (**9**).

Peak at δ 7.26 is CHCl_3_, and peak at δ 7.0 is unknown impurity.

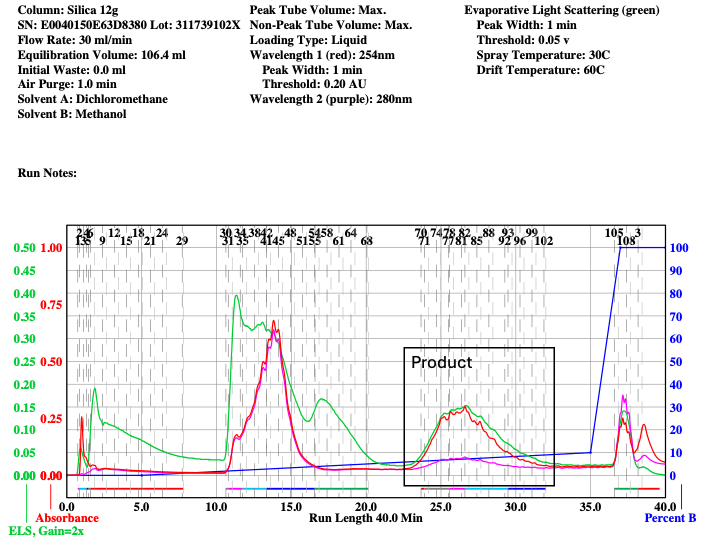

Figure S16. ISCO chromatogram for oxaphosphinanes (**10**).

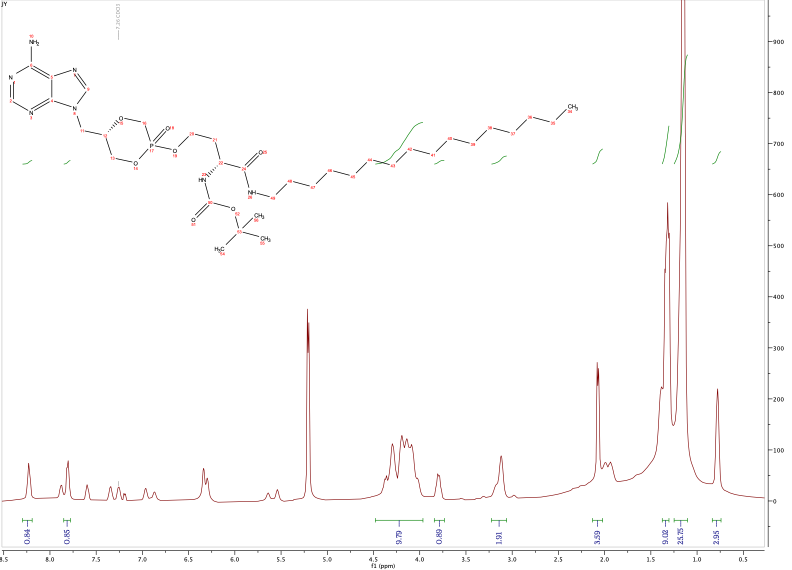

Figure S17. ^1^H NMR (600 MHz, CDCl_3_) spectrum of oxaphosphinanes (**10**).

Trace peaks in the aromatic region are from side products which were removed via flash chromatography in the next step.

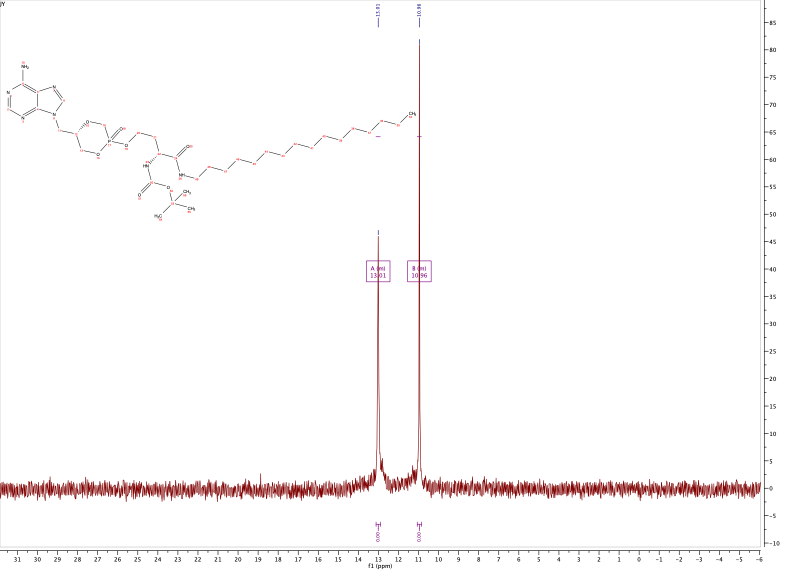

Figure S18. ^31^P NMR (243 MHz, CDCl_3_) spectrum of oxaphosphinanes (**10**).

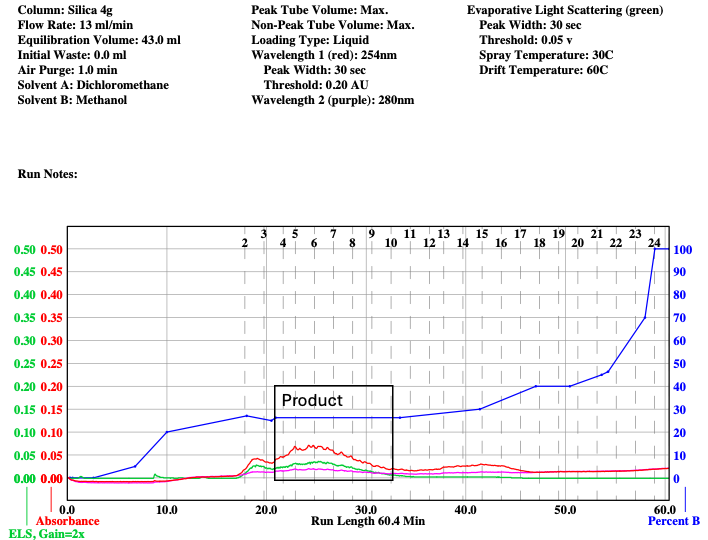

Figure S19. ISCO chromatogram for phosphonate (**11**).

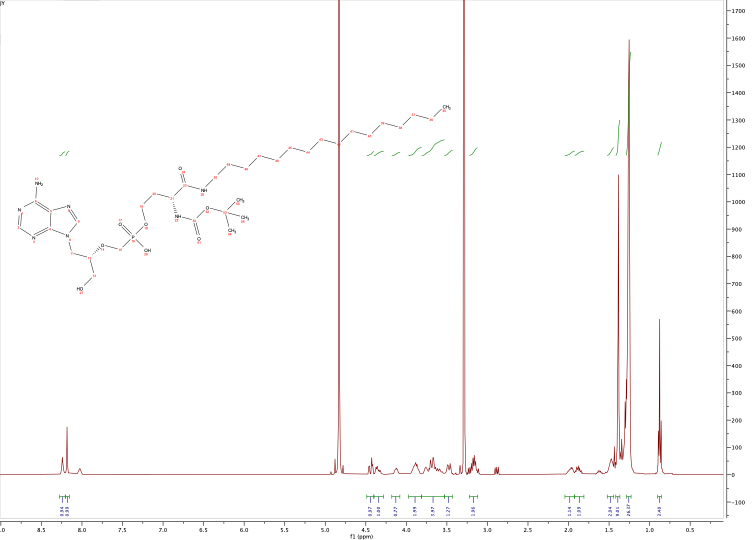

Figure S20. ^1^H NMR (400 MHz, CD_3_OD) spectrum of phosphonate (**11**).

Contaminant at δ 8.0 was removed in the subsequent step.

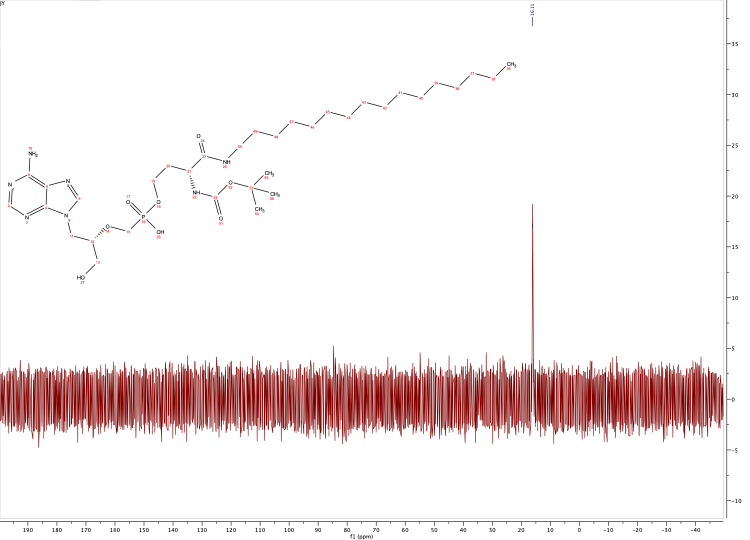

Figure S21. ^31^P NMR (162 MHz, CD_3_OD) spectrum of phosphonate (**11**).

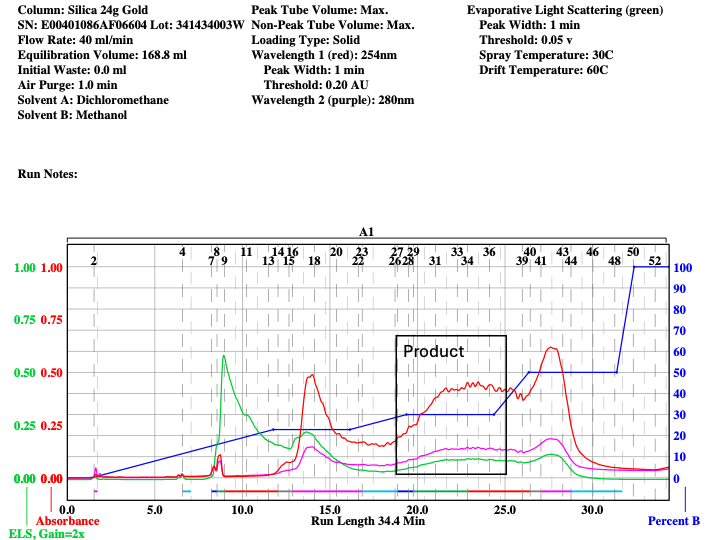

Figure S22. ISCO chromatogram for USC-093D (**12**).

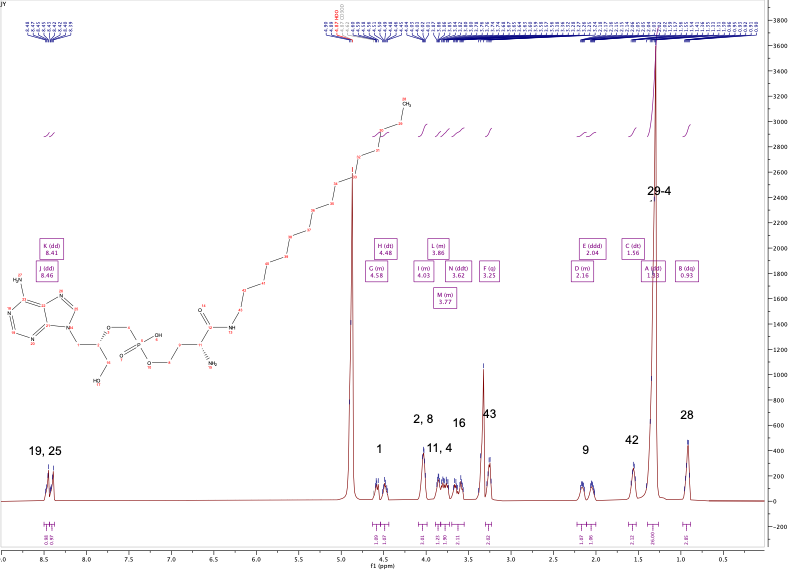

Figure S23. ^1^H NMR (600 MHz, CD_3_OD) spectrum of USC-093D (**12**).

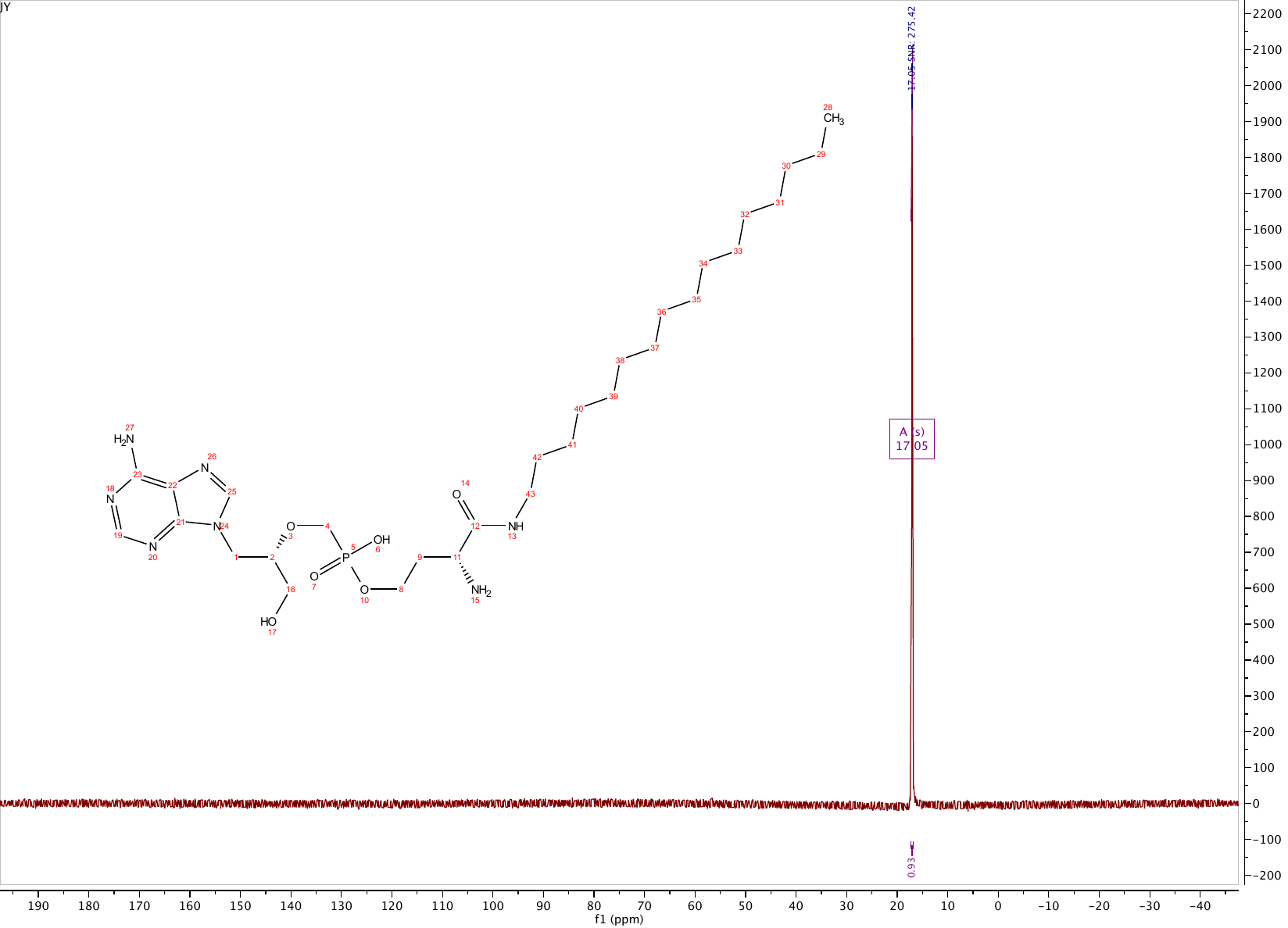

Figure S24. ^31^P NMR (243 MHz, CD_3_OD) spectrum of USC-093D (**12**).

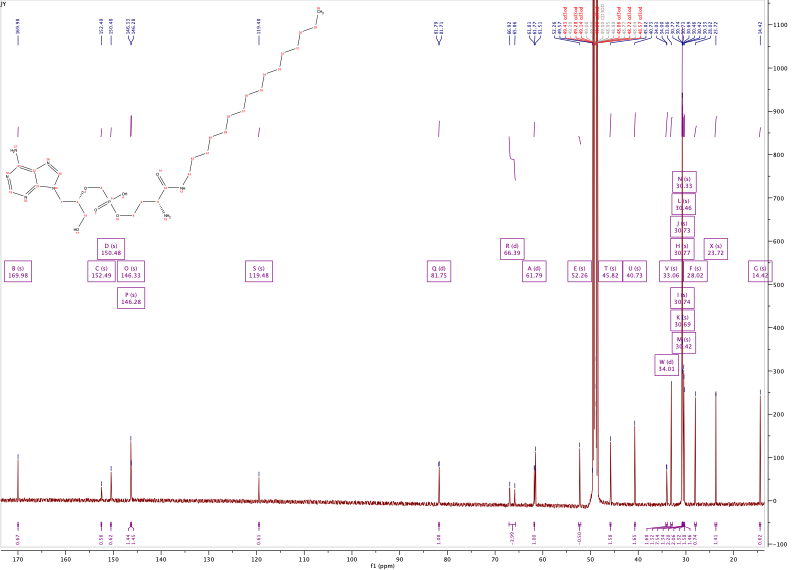

Figure S25. ^13^C NMR (151 MHz, CD_3_OD) spectrum of USC-093D (**12**).

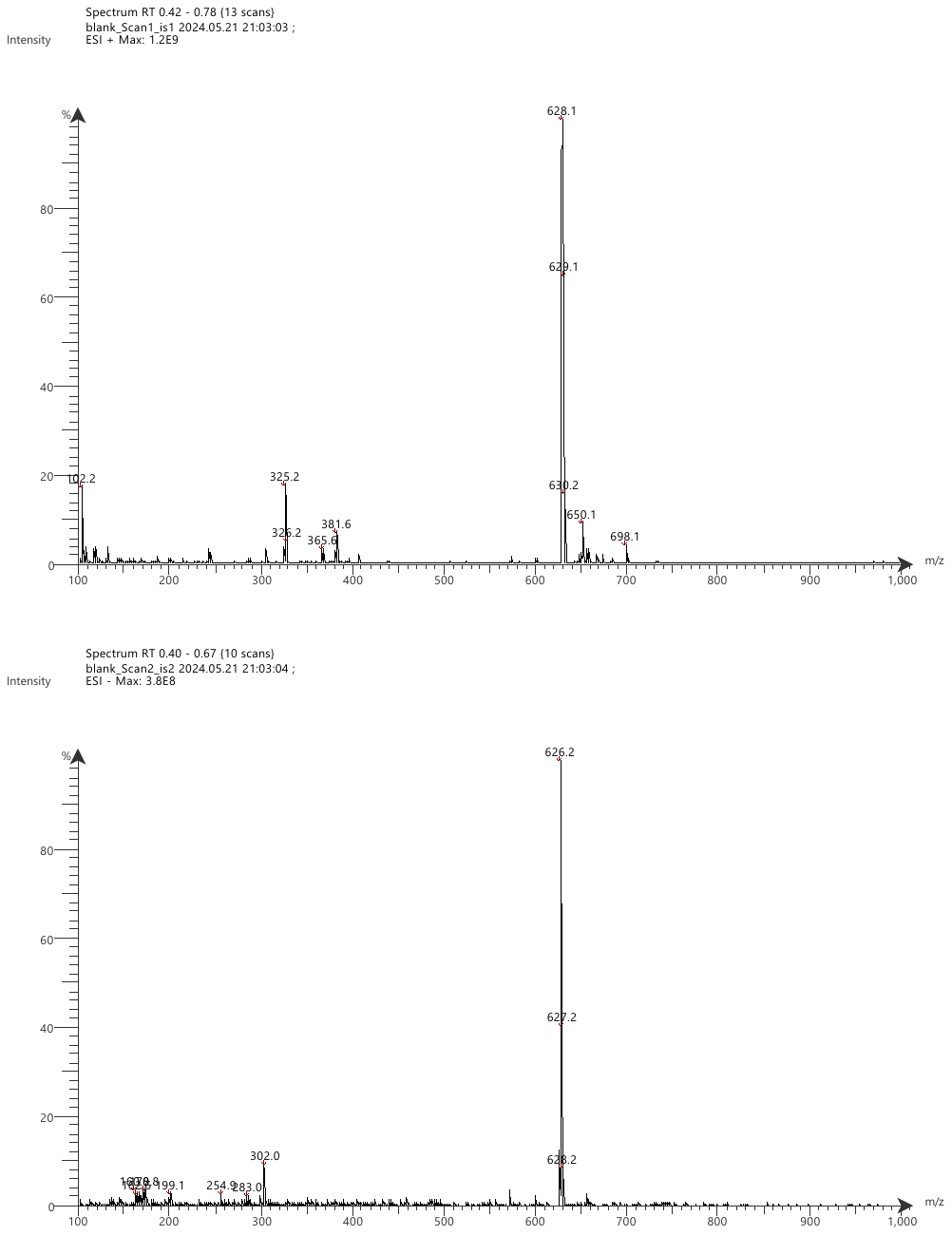

Figure S26. MS spectrum of USC-093D (**12**) in negative mode.

[M-H]^-^: C_29_H_53_N_7_O_6_P^-^; calcd 626.39, found 626.2 – Advion.

| ID | NA | K | CL | GLU | BUN | CREAT | CA | PHOS | TP | ALB | GLOB | CHOL | TRIG | AST | ALT | GGT | TBILI | ALP | CK | AMY | LIP | CO2 | ANION | OSMO | CALP |
| --- | --- | --- | --- | --- | --- | --- | --- | --- | --- | --- | --- | --- | --- | --- | --- | --- | --- | --- | --- | --- | --- | --- | --- | --- | --- |
| 1 | 144.0 | 6.2 | 94.0 | 84 | 19.1 | 0.33 | 13.1 | 9.2 | 5.1 | 3.31 | 1.8 | 83 | 167 | 363.4 | 127.1 | 0.3 | 0.01 | 108.5 | 2956 | 1780 | 109 | 37.4 | 18.8 | 299.9 | 3.0 |
| 2 | 139.0 | 7.0 | 92.0 | 173 | 20.0 | 0.31 | 13.9 | 10.8 | 5.2 | 3.33 | 1.9 | 113 | 172 | 53.6 | 66.0 | 0.7 | 0.02 | 129.5 | 319 | 1102 | 36 | 36.0 | 18.0 | 297.3 | 6.0 |
| 3 | 142.0 | 6.7 | 95.0 | 116 | 19.1 | 0.28 | 12.7 | 9.0 | 5.4 | 3.42 | 2.0 | 99 | 187 | 184.5 | 95.1 | 0.2 | 0.03 | 137.0 | 1346 | 1750 | 37 | 34.6 | 19.1 | 298.8 | 5.0 |
| 4 | 137.0 | 7.7 | 89.0 | 214 | 20.5 | 0.25 | 14.5 | 10.4 | 5.2 | 3.28 | 1.9 | 87 | 144 | 36.6 | 52.0 | 0.8 | 0.01 | 105.5 | 223 | 2068 | 32 | 37.7 | 18.0 | 297.4 | 7.0 |
| 5 | 145.0 | 7.0 | 94.0 | 85 | 20.0 | 0.35 | 13.0 | 10.1 | 5.4 | 3.49 | 1.9 | 111 | 178 | 160.7 | 53.0 | 0.3 | 0.05 | 132.1 | 1299 | 1150 | 39 | 39.3 | 18.7 | 303.6 | 5.0 |
| 6 | 138.0 | 7.9 | 92.0 | 248 | 19.8 | 0.29 | 14.4 | 11.1 | 5.4 | 3.52 | 1.9 | 132 | 193 | 65.6 | 102.7 | 1.3 | 0.05 | 146.2 | 381 | 1636 | 36 | 33.0 | 20.9 | 301.2 | 10.0 |
| 7 | 142.0 | 7.6 | 96.0 | 183 | 18.7 | 0.27 | 13.4 | 9.2 | 5.5 | 3.46 | 2.0 | 117 | 165 | 140.7 | 28.3 | 0.5 | 0.02 | 128.5 | 952 | 1544 | 40 | 30.9 | 22.7 | 304.1 | 4.0 |
| 8 | 141.0 | 8.0 | 91.0 | 324 | 21.0 | 0.34 | 15.3 | 10.2 | 5.8 | 3.67 | 2.1 | 129 | 196 | 63.2 | 58.5 | 0.9 | 0.06 | 134.4 | 256 | 1962 | 36 | 33.1 | 24.9 | 311.6 | 8.0 |
| 9 | 140.0 | 8.5 | 92.0 | 345 | 17.3 | 0.26 | 14.0 | 10.2 | 5.3 | 3.53 | 1.8 | 105 | 157 | 57.5 | 81.6 | 0.8 | 0.03 | 129.7 | 264 | 1592 | 36 | 33.0 | 23.5 | 310.6 | 7.0 |
| 10 | 142.0 | 7.6 | 93.0 | 147 | 20.7 | 0.28 | 12.8 | 9.3 | 5.6 | 3.54 | 2.1 | 110 | 165 | 398.0 | 381.1 | 0.1 | 0.03 | 146.6 | 1968 | 1478 | 40 | 33.4 | 23.2 | 302.8 | 3.0 |
| 11 | 144.0 | 6.6 | 94.0 | 108 | 18.9 | 0.31 | 13.4 | 8.8 | 5.8 | 3.78 | 2.0 | 101 | 193 | 206.0 | 60.4 | 0.4 | 0.01 | 128.9 | 1464 | 2022 | 35 | 32.4 | 24.2 | 301.9 | 6.0 |
| 12 | 142.0 | 7.5 | 93.0 | 246 | 19.1 | 0.33 | 14.2 | 10.3 | 5.5 | 3.55 | 2.0 | 111 | 194 | 49.6 | 84.5 | 0.4 | 0.03 | 146.8 | 265 | 1749 | 104 | 34.0 | 22.5 | 307.6 | 9.0 |
| 13 | 137.0 | 7.9 | 91.0 | 209 | 20.6 | 0.33 | 13.6 | 9.7 | 5.4 | 3.42 | 2.0 | 93 | 177 | 459.3 | 203.8 | 0.4 | 0.04 | 113.6 | 3993 | 1728 | 151 | 35.1 | 18.8 | 297.5 | 4.0 |
| 14 | 138.0 | 8.3 | 92.0 | 237 | 17.6 | 0.34 | 13.9 | 10.8 | 4.7 | 3.23 | 1.5 | 81 | 147 | 41.7 | 49.2 | 1.0 | 0.05 | 137.9 | 140 | 1489 | 38 | 33.2 | 21.1 | 300.6 | 8.0 |
| 15 | 142.0 | 8.3 | 96.0 | 229 | 17.8 | 0.32 | 14.1 | 10.2 | 5.2 | 3.54 | 1.7 | 89 | 141 | 60.0 | 52.8 | 0.8 | 0.03 | 108.3 | 184 | 1170 | 53 | 34.8 | 19.5 | 307.6 | 8.0 |
| 16 | 141.0 | 7.3 | 94.0 | 143 | 17.6 | 0.24 | 13.7 | 9.2 | 5.5 | 3.61 | 1.9 | 95 | 180 | 167.9 | 35.5 | 0.2 | 0.02 | 151.0 | 874 | 2136 | 37 | 33.3 | 21.0 | 299.1 | 7.0 |
| 17 | 144.0 | 7.9 | 95.0 | 263 | 19.2 | 0.29 | 14.8 | 10.5 | 5.5 | 3.39 | 2.1 | 92 | 178 | 69.7 | 99.1 | 0.6 | 0.04 | 113.1 | 219 | 1737 | 47 | 33.3 | 23.6 | 313.0 | 9.0 |
| 18 | 143.0 | 8.3 | 95.0 | 207 | 20.1 | 0.29 | 14.0 | 10.7 | 5.4 | 3.62 | 1.8 | 97 | 165 | 75.1 | 95.2 | 0.7 | 0.03 | 135.8 | 275 | 1685 | 40 | 34.0 | 22.3 | 309.1 | 8.0 |
| 19 | 147.0 | 7.1 | 114.0 | 142 | 14.3 | 0.26 | 14.2 | 8.1 | 5.4 | 3.16 | 2.2 | 123 | 162 | 138.8 | 41.7 | 0.6 | 0.05 | 126.8 | 902 | 2470 | 50 | 22.7 | 17.4 | 308.6 | 10.0 |
| 20 | 148.0 | 6.9 | 109.0 | 162 | 12.9 | 0.31 | 14.0 | 8.5 | 5.8 | 3.69 | 2.1 | 102 | 169 | 137.5 | 50.1 | 0.9 | 0.04 | 177.5 | 756 | 2251 | 35 | 27.6 | 18.3 | 310.7 | 10.0 |
| 21 | 141.0 | 7.2 | 105.0 | 178 | 13.6 | 0.29 | 14.2 | 7.4 | 5.4 | 3.28 | 2.1 | 95 | 155 | 54.8 | 40.3 | 0.9 | 0.04 | 138.3 | 233 | 1967 | 46 | 27.4 | 15.8 | 299.4 | 10.0 |
| 22 | 143.0 | 7.1 | 104.0 | 128 | 13.5 | 0.29 | 12.6 | 7.4 | 5.3 | 3.30 | 2.0 | 89 | 125 | 214.4 | 52.7 | 0.1 | 0.02 | 168.0 | 1736 | 1896 | 38 | 29.5 | 16.6 | 300.1 | 2.0 |
| 23 | 144.0 | 7.3 | 109.0 | 175 | 16.7 | 0.26 | 14.1 | 8.1 | 5.4 | 3.28 | 2.1 | 100 | 123 | 109.1 | 68.9 | 0.3 | 0.05 | 153.9 | 796 | 2134 | 48 | 25.6 | 16.7 | 306.1 | 12.0 |
| 24 | 141.0 | 7.5 | 106.0 | 204 | 18.6 | 0.26 | 13.7 | 8.6 | 5.5 | 3.36 | 2.1 | 104 | 95 | 88.0 | 64.4 | 1.5 | 0.07 | 165.6 | 270 | 2220 | 35 | 22.2 | 20.3 | 303.2 | 13.0 |
| 25 | 147.0 | 7.3 | 97.0 | 149 | 21.8 | 0.30 | 13.9 | 10.2 | 5.6 | 3.60 | 2.0 | 119 | 166 | 210.9 | 58.7 | 0.8 | 0.02 | 151.7 | 1673 | 1703 | 53 | 30.2 | 27.1 | 312.1 | 8.0 |
| 26 | 171.0 | 12.0 | 120.0 | 96 | 24.0 | 0.42 | 15.3 | 13.8 | 6.6 | QNS | QNS | 138 | 201 | 544.5 | 207.0 | QNS | QNS | 157.2 | 6021 | 1812 | 249 | QNS | QNS | 363.3 | 21.0 |
| 27 | 143.0 | 8.1 | 95.0 | 244 | 19.6 | 0.28 | 14.3 | 10.9 | 5.6 | 3.54 | 2.1 | 121 | 198 | 73.6 | 125.5 | 0.9 | 0.02 | 111.8 | 344 | 1651 | 39 | 30.5 | 25.6 | 310.6 | 8.0 |
| 28 | 146.0 | 8.2 | 97.0 | 210 | 21.4 | 0.31 | 14.1 | 10.7 | 5.5 | 3.62 | 1.9 | 111 | 168 | 44.7 | 44.5 | 1.0 | 0.03 | 130.1 | 297 | 1653 | 50 | 33.2 | 24.0 | 315.1 | 10.0 |
| 29 | 146.0 | 8.1 | 95.0 | 289 | 18.6 | 0.27 | 15.1 | 10.4 | 5.6 | 3.61 | 2.0 | 106 | 177 | 75.7 | 70.1 | 0.6 | 0.02 | 140.9 | 564 | 1881 | 39 | 32.4 | 26.7 | 318.3 | 9.0 |
| 30 | 146.0 | 8.3 | 95.0 | 234 | 18.3 | 0.28 | 15.0 | 10.6 | 5.9 | 3.75 | 2.2 | 117 | 175 | 67.7 | 113.4 | 0.4 | 0.02 | 158.3 | 278 | 1847 | 41 | 31.8 | 27.5 | 315.5 | 10.0 |
| 31 | 147.0 | 6.7 | 96.0 | 202 | 19.1 | 0.28 | 14.7 | 10.4 | 5.7 | 3.71 | 2.0 | 124 | 196 | 55.8 | 61.6 | 0.9 | 0.04 | 150.9 | 275 | 1735 | 42 | 31.0 | 26.7 | 312.9 | 10.0 |
| 32 | 145.0 | 7.9 | 97.0 | 186 | 20.7 | 0.38 | 14.3 | 10.8 | 5.3 | 3.46 | 1.8 | 104 | 178 | 60.1 | 56.9 | 0.9 | 0.03 | 147.6 | 227 | 1644 | 36 | 31.5 | 24.4 | 311.1 | 9.0 |
| 33 | 144.0 | 8.0 | 94.0 | 246 | 22.6 | 0.26 | 14.7 | 10.1 | 5.6 | 3.57 | 2.0 | 106 | 201 | 48.5 | 75.3 | 1.2 | 0.05 | 154.5 | 179 | 1982 | 37 | 30.6 | 27.4 | 313.5 | 11.0 |
| 34 | 150.0 | 7.7 | 98.0 | 149 | 17.3 | 0.25 | 13.7 | QNS | 6.0 | QNS | QNS | QNS | 197 | 112.8 | 70.9 | QNS | QNS | 125.1 | 662 | 2138 | QNS | QNS | QNS | 316.8 | QNS |
| 35 | 149.0 | 8.1 | 95.0 | QNS | 23.0 | 0.30 | QNS | QNS | 6.0 | QNS | QNS | QNS | QNS | 37.2 | 58.9 | QNS | QNS | 155.7 | QNS | 1779 | QNS | QNS | QNS | QNS | QNS |
| 36 | 147.0 | 8.4 | 97.0 | 240 | 22.3 | 0.32 | 14.9 | 10.0 | 6.1 | 4.02 | 2.1 | 109 | 164 | 54.7 | 59.8 | 0.7 | 0.04 | 116.2 | 299 | 2045 | 41 | 30.1 | 28.3 | 319.3 | 10.0 |
| 37 | 145.0 | 7.7 | 102.0 | 209 | 16.6 | 0.31 | 14.2 | 9.6 | 5.3 | 3.32 | 2.0 | 107 | 171 | 51.7 | 47.4 | 0.6 | 0.03 | 111.2 | 184 | 1655 | 41 | 29.5 | 21.2 | 310.6 | 9.0 |
| 38 | 145.0 | 7.7 | 106.0 | 177 | 15.5 | 0.31 | 13.8 | 8.9 | 5.2 | 3.32 | 1.9 | 88 | 144 | 41.5 | 34.4 | 1.0 | 0.27 | 149.2 | 201 | 1781 | 36 | 26.6 | 20.1 | 308.4 | 9.0 |
| 39 | 143.0 | 7.9 | 103.0 | 265 | 15.4 | 0.32 | 14.9 | 8.5 | 5.7 | 3.74 | 2.0 | 94 | 182 | 51.3 | 40.6 | 1.3 | 0.08 | 148.6 | 184 | 1936 | 34 | 27.0 | 20.9 | 309.9 | 12.0 |
| 40 | 148.0 | 7.3 | 101.0 | 102 | 18.7 | 0.30 | 12.6 | 8.9 | 5.3 | 3.42 | 1.9 | 91 | 167 | 89.1 | 41.1 | 1.2 | 0.01 | 147.6 | 617 | 1554 | 37 | 33.3 | 21.0 | 310.2 | 10.0 |
| 41 | 144.0 | 7.7 | 97.0 | 311 | 19.0 | 0.28 | 14.3 | 9.7 | 5.4 | 3.41 | 2.0 | 87 | 170 | 131.5 | 92.2 | 0.6 | 0.01 | 120.9 | 1197 | 1936 | 43 | 27.3 | 27.4 | 315.2 | 1.0 |
| 42 | 144.0 | 8.2 | 97.0 | 218 | 19.1 | 0.34 | 14.5 | 10.1 | 5.3 | 3.45 | 1.9 | 88 | 184 | 48.0 | 43.8 | 0.7 | 0.02 | 136.8 | 217 | 1758 | 38 | 29.5 | 25.7 | 311.0 | 6.0 |

Table S1. Serum Chemistry.
